# Mapping polyclonal antibody responses in non-human primates vaccinated with HIV Env trimer subunit vaccines

**DOI:** 10.1101/833715

**Authors:** Bartek Nogal, Matteo Bianchi, Christopher A. Cottrell, Robert N. Kirchdoerfer, Leigh M. Sewall, Hannah L. Turner, Fangzhu Zhao, Devin Sok, Dennis R. Burton, Lars Hangartner, Andrew B. Ward

## Abstract

Rational immunogen design aims to focus antibody responses to vulnerable sites on the primary antigens. Given the size of these antigens there is however potential for eliciting unwanted, off-target responses. Here, we used our electron microscopy polyclonal epitope mapping approach to describe the antibody specificities elicited by immunization of non-human primates with soluble HIV envelope trimers and subsequent repeated viral challenge. An increased diversity of epitopes recognized, and the approach angle by which these antibodies bound, constituted a hallmark of the humoral response in most protected animals. We also show that fusion peptide-specific antibodies are responsible for some neutralization breadth. Moreover, cryoEM analysis of a fully-protected animal revealed a high degree of clonality within a subset of putatively neutralizing antibodies, enabling a detailed molecular description of the antibody paratope. Our results provide important insights into the immune response against a vaccine candidate that entered into clinical trials earlier this year.

## INTRODUCTION

Despite marked progress toward HIV management with effective anti-retroviral therapy (ART) options, HIV and acquired immunodeficiency syndrome (AIDS) remain major global health challenges as we approach the end of the second decade of the 21^st^ century. While ART and other prevention options have reduced the overall incidence of HIV/AIDS, (Cohen et al., 2011), nearly a million individuals are newly infected each year. Thus, a vaccine that protects from infection remains the most robust approach in eradicating HIV across populations. However, due in no small part to the high rate of mutation of HIV and the associated mastery of disguise by a dense and adaptable glycan shield, efforts toward developing an immunogen that would impart protection from global HIV subtypes have thus far failed. While correlates and mechanisms of vaccine protection are still being confirmed in clinical trials, the passive transfer of neutralizing antibodies in rhesus macaques have shown to reproducibly protect against simian-human immunodeficiency virus (SHIV) challenge (Ahmed et al., 2017; Burton and Hangartner, 2016; Lu et al., 2016). Thus, broadly neutralizing antibodies (bnAbs) and the associated epitopes that they engage on the HIV envelope protein (Env) have helped to inform HIV rational vaccine design, with the ultimate goal of bnAb elicitation and thus protection from global HIV subtypes via immunization.

HIV bnAbs are present in a minority of HIV-1 infected individuals and are present at relatively rare frequencies in the overall humoral response to infection (Cohen and Frahm, 2017). The epitopes targeted by bnAbs include the CD4-binding site (CD4bs), the variable region 1 and 2 glycan site (V1/V2-glycan), the N332 supersite, the gp120-gp41 interface, the silent face (VRC-PG05, (Zhou et al., 2018)), and the membrane proximal external region (MPER) (Burton and Hangartner, 2016). BnAbs generally have atypical features compared to the normal antibody repertoire including unusually long heavy chain complementary determining region 3 (CDRH3) or short CDRL3s, auto- and poly-reactivity, and/or high levels of somatic hypermutation (SHM) (West et al., 2014; Yu and Guan, 2014). Moreover, many germline-reverted bnAbs have little or no affinity for the HIV envelope trimer (Escolano et al., 2016; Prabakaran et al., 2014). Thus, elicitation of bnAbs within the practical framework of a prime-boost vaccination approach is not trivial. To date, human clinical trials have shown no or limited impact, with only one, based on gp120, showing limited efficacy (Haynes et al., 2016; Hsu and O’Connell, 2017; Rerks-Ngarm et al., 2009). Recent animal studies using novel immunogens designed to present a more native structure and antigenic surface of the Env trimer are however defining a new and potentially promising path toward eliciting more productive immune responses (Jardine et al., 2013).

One such protein subunit immunogen is the BG505 SOSIP.664 gp140 trimer (https://clinicaltrials.gov/ct2/show/NCT03699241), a mimic of the natural Env spike protein, that is stabilized in a prefusion conformation, which is a state recognized by broadly neutralizing but not non-neutralizing Abs (Boris D Juelg, 2018; Chuang et al., 2017; Julien et al., 2013; McElrath, Julie, Omu Anzala, 2018; Pugach et al., 2015). This immunogen recapitulates the quaternary structural features and surface glycans that Abs must learn to navigate as they mature into potent and broad responses via somatic hypermutation (SHM). As this vaccine candidate enters human clinical trials (McElrath, Julie, Omu Anzala, 2018), an examination of elicited antibody response in an animal model that more closely approximates the complexity of the human immune system is timely. Pauthner et al. recently reported success in eliciting protective tier-2 virus neutralization responses by BG505 SOSOP.664 immunization in rhesus macaques (Pauthner *et al*., 2017). Immunized non-human primates (NHP) were grouped into “high-titer” or “low-titer” animals, depending on the ability of their serum antibodies to neutralize BG505 pseudo-virus. Both groups, matched with non-immunized controls, were then repetitively challenged with a chimeric simian-human immunodeficiency virus (SHIV) expressing the autologous BG505 Env (SHIV_BG505_) (Pauthner et al., 2018). Since the variable levels of both neutralization titers and protection from challenge observed in these macaques may hold important clues as to the potentially productive and unproductive immune system paths that may be evoked by the native-like SOSIP trimer, we decided to perform an analysis of epitopes recognized in the animals from these groups, and to compare them to the antibody response elicited following infection of non-immunized animals.

Despite its native antigenic profile, the BG505 SOSIP.664 trimer exposes surfaces such as the base of the trimer (BOT) that are not accessible on the full-length, membrane inserted Env protein, and that have been shown to be very immunogenic in the rabbits (Bianchi et al., 2018). While humoral responses against the BOT are irrelevant, the responses to the other regions of the trimer have the ability to elicit protective tier-2 neutralizing antibodies. However, these responses typically comprise variable epitopes that are strain-specific with little to no potential for developing breadth. Further, these unwanted responses may be competing with elicitation of more desirable bnAb. We currently only have a limited understanding of the diversity in the responses to SOSIP trimers which has been derived primarily from rabbit immunization studies (Bianchi et al., 2018; Klasse et al., 2018; Mccoy et al., 2016; Pauthner et al., 2017). To investigate the specificity of polyclonal antibody responses, we recently developed an electron microscopy-based polyclonal epitope mapping technique, whereby total serum IgG is isolated, digested in fragments antigen binding (Fabs) and complexed with Env SOSIP.664 trimers before they are imaged using negative-stain or cryoEM. This technique provides a visual snapshot of the specificities present in polyclonal antibody preparations of an individual at a given time. We have thus far used electron microscopy polyclonal epitope mapping (EMPEM) to define epitopes recognized by sera of BG505 SOSIP.664 immunized rabbits. The responses were shown to be quite similar between individual animals and to recognize a relatively narrow range of epitopes (Bianchi et al., 2018).

Here, we report EMPEM analysis of NHP polyclonal responses to BG505-based immunogens described in the Pauthner et al. studies (Pauthner et al., 2017, 2018). We found that high-titer animals recognized a greater diversity of epitopes than lower titer animals. Our analyses also identified several strain-specific responses that might sterically block access to bnAb epitopes. Further, by using heterologous SOSIP trimers, we were able to map the epitope of cross-reactive antibodies that were detected in several immunized animals. It was found to be located at the gp120/gp41 interface region that includes the fusion peptide. Finally, our ∼3.8 Å resolution cryoEM reconstruction of Fabs from a fully protected animal complexed to SOSIP trimers illustrated that one of the specificities, V1/V3, was likely a clonal response. This structure also revealed that the V1/V3 loop antibody that resembles N332 supersite bnAbs at low resolution primarily interacts with the V1 loop at high resolution, and that their binding would sterically compete with N332 supersite-specific bnAbs. In total, our imaging results provide many new insights for Env trimer immunogen redesign including both positive and negative engineering opportunities.

## RESULTS

### Humoral Responses in Naïve Macaques After SHIV_BG505_ Infection

The SHIV_BG505_ virus challenge study enrolled 12 immunized and 6 naïve control animals (Pauthner et al., 2018). As previously described, the challenging virus contained a S375Y mutation that confers replication capacity in rhesus macaques and results in AIDS-like pathology (Pauthner et al., 2018). Animals received up to 12 intra rectal challenges at weekly intervals with a break after 6 challenges for those animals not infected early. 5/6 naïve animals became infected after the first challenge, and one after the second. In the low-titer NHP group, 2/6 animals became infected after the first challenge, and 4/6 after the second. Finally, in the high-titer NHP group 1/6 animals became infected after the first challenge, 1/6 after the sixth, 1/6 after the tenth, 1/6 after the twelfth, and 2/6 never became infected (Pauthner et al., 2018).

Using EMPEM, we first analyzed the antibody responses in week 20 post challenge (p.c.) sera (i.e. 20 weeks post first challenge) of the six non-immunized animals by forming complexes with BG505 SOSIP.664v5.2 Env glycoprotein trimers that were sequence matched to immunogen and the Env of the SHIV_BG505_ challenge virus. Figure 1 summarizes the EMPEM results wherein each dataset was reconstructed into a global 3D average, and then subjected to subsequent 3D classification that sorted data into unique classes (Figure S1). Two of the six serum samples analyzed, BZ15 and BZ31, were not amenable to 3D reconstruction. The epitopes recognized in these animals however could be discerned from the 2D class averages when compared to known references (Figure S1D and S1E). One animal, CB00, showed no detectable binding to the BG505 trimer probe (Figure S1F), yet this was not unexpected since this animal showed waning BG505 S375Y pseudo-virus neutralizing titers 12 weeks p.c. (Pauthner et al., 2018), suggesting viral evolution away from the infecting virus and concomitant antibody evolution. The remaining three animals exhibited antibody responses in and around the N289/S241 glycan hole region (GH-1/GH-2) that is a highly immunogenic epitope in rabbits (McCoy *et al*., 2016; Bianchi *et al*., 2018; Klasse *et al*., 2018), and the C3/V5 region of gp120 that we have previously described as the “C3/T465 epitope” in NHPs. Thus, considering a sample of only six animals and a time span of just 20 weeks under viral challenge pressure, the infected control macaques exhibited a relatively narrow repertoire of antibody epitope responses to the founding virus, with five different 3D Fab phenotypes detected against two regions of Env (Figure 1B and 1C). Notably, and not unexpectedly, virus challenges did not induce any antibodies that bound to the base of the trimer (Figure 1), which in rabbits is typically immunodominant on the soluble trimer and may limit elicitation of neutralizing immune responses (McCoy *et al*., 2016; Bianchi *et al*., 2018).

**Figure 1.**
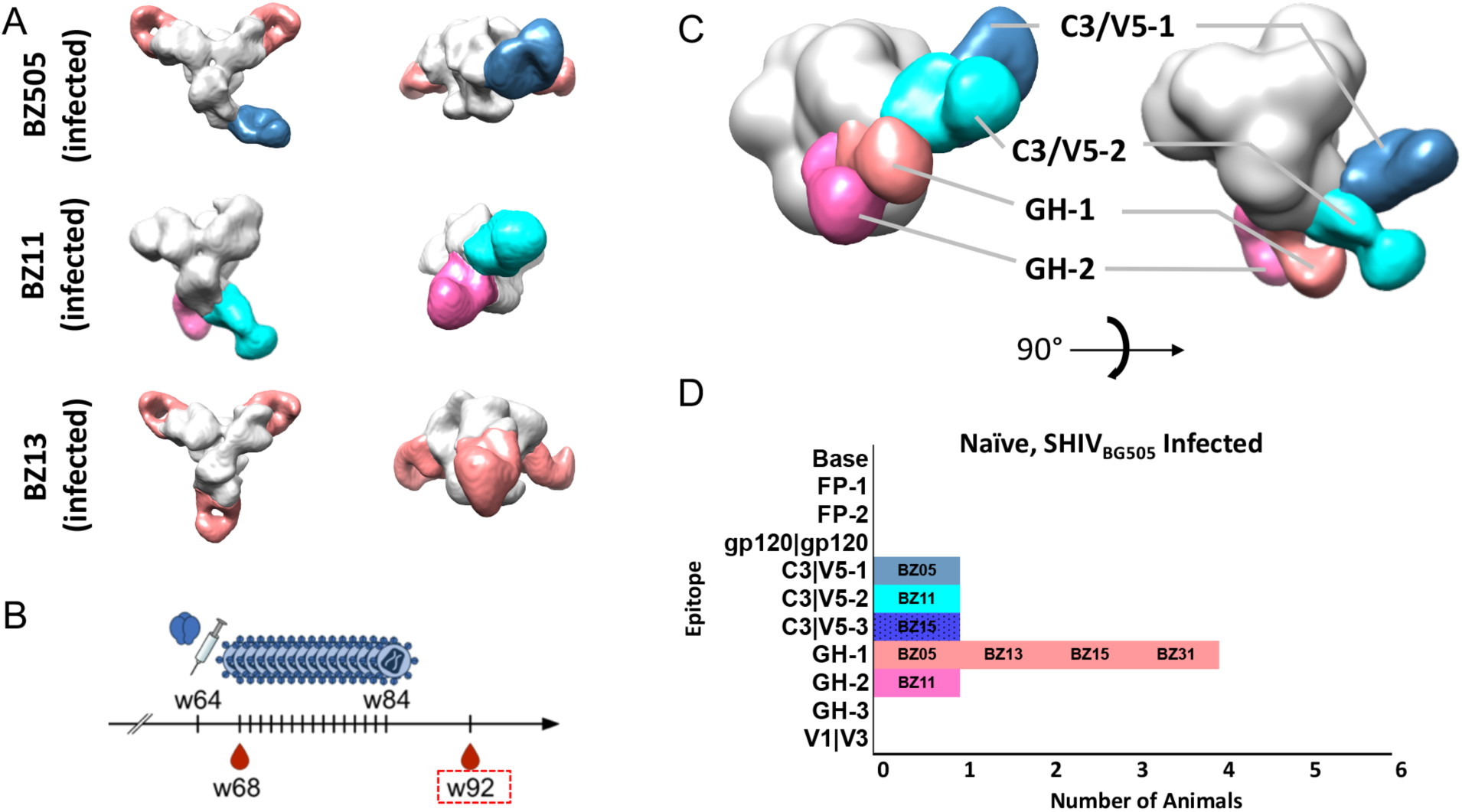
Polyclonal antibody responses of naïve macaques infected by SHIV_BG505_ challenge. (A) Segmented 3D reconstructions of polyclonal antibody-BG505 trimer complexes from animals BZ05, BZ11, and BZ13. (B) Schedule of SHIV_BG505_ challenges. The red box on the timeline indicates the week from which sera were used for EM imaging. (C) Composite 3D reconstruction demonstrating the antigenic map of all epitopes observed in SHIV_BG505_ infected macaques (see Figure S2 for 3D classification and 2D class averages of animals BZ31, BZ15, and CB00). (D) Bar graph summary of the epitopes targeted by SHIV_B505_ infected macaques (BZ15 phenotype represented in bar graph by 2D classification only).

### Low nAb Titer Vaccinated Animals Show Increased Diversity of Epitopes Compared to Infected Naïve Controls

The next group of low-titered animals that we examined was subjected to a total of 4 identical immunizations with BG505 SOSIP.664 variants, including a booster injection 4 weeks prior to the first SHIV_BG505_ challenge. We performed EMPEM analysis on these animals 4 weeks after this last booster, hence immediately prior to the first virus challenge (i.e. week 68). These samples showed an overlapping set of epitopes but with increased diversity relative to the infected naïve controls (Figures 1 and 2). The primary differences were that immunized animals also mounted an antibody response to the fusion peptide region, the gp120/gp120 interface, and the base of the trimer. Despite the diversity of epitopes recognized, all animals from this low-titer group became infected within the first two virus challenges (Pauthner et al., 2018). Interestingly, and perhaps driven by the presence of BG505 SOSIP vaccination-induced antibodies, these animals had reduced peak viral loads compared to the immunogen-naïve controls (Pauthner et al., 2018). Second to base binding, which is present in all BG505 SOSIP immunized animals (McCoy *et al*., 2016; Bianchi *et al*., 2018), the most common epitopes targeted in the low-titer animals were the glycan hole 1 (GH-1) epitope, which was also the most frequently recognized epitope among the unimmunized controls, and the fusion peptide (FP; Figures 1B, 1C and 2G). There was no obvious correlation between epitopes recognized and protection or viral titer (Pauthner et al., 2018). Thus, despite the presence of a diversity of vaccine-induced antibodies, this group of animals was not protected from the virus challenge. We note that none of the animals in this group had neutralizing antibody titers above the ∼1:500 threshold determined as the main correlate of protection (Pauthner et al., 2018).

**Figure 2.**
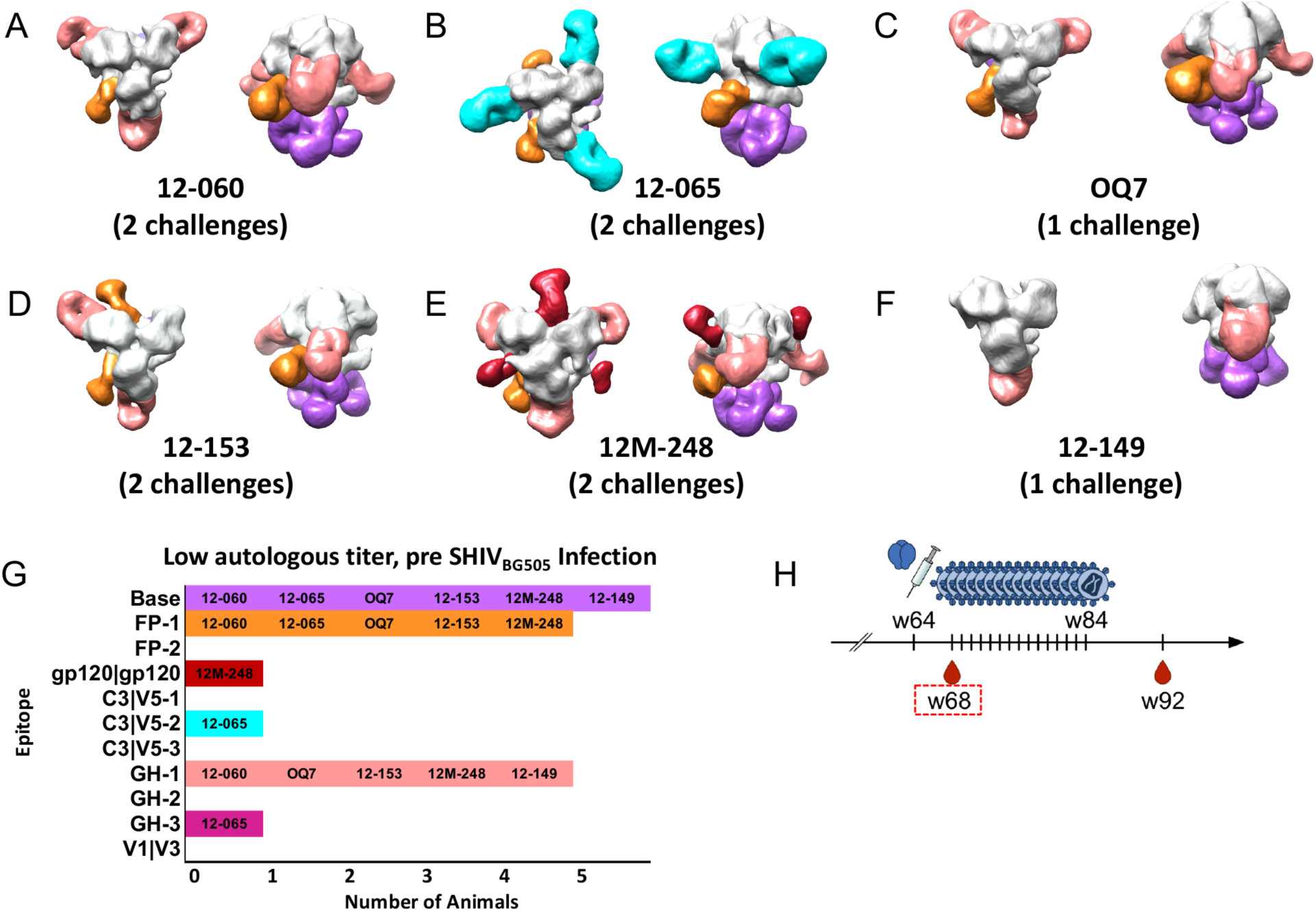
Polyclonal antibody responses of low autologous neutralizing titer macaques immediately before SHIV_BG505_ challenge. Segmented 3D reconstructions of (A) 12-060, (B) 12-065, OQ7, (D) 12-153, (E) 12M-248, and (F) 12-149 and (G) Bar graph summary of the epitopes targeted by low autologous titer macaques. The number in parentheses indicates the number of SHIV_BG505_ challenges it took for the animal to become infected (see Figure S2 for additional 3D phenotypes). (H) The red box on timeline indicates the week from which sera were used for EM imaging. Week 68 is four weeks after the last boost.

### High nAb Titer Vaccinated Animals Show the Most Diverse Humoral Responses

Relative to the infected naïve and the low-titer immunized macaques, the high-titer animals exhibited overlapping yet further diversified responses; their antibody response not only differed in the number and types of epitopes recognized, but also in the apparent angles of approach for similar epitope clusters (Figures 1, 2, and 3). This higher diversity probably also gave rise to an increased Fab occupancy in the complexes observed in these animals. Figure 3 shows the results of the EMPEM analysis of week 68 serum samples from the high-titer macaque group that showed significant protection from infection and reduced peak viral loads relative to naïve animals (Pauthner et al., 2017, 2018). Notably, two animals, 12-046 and 4O9, had remained uninfected after twelve SHIV_BG505_challenges (Pauthner et al., 2018).

**Figure 3.**
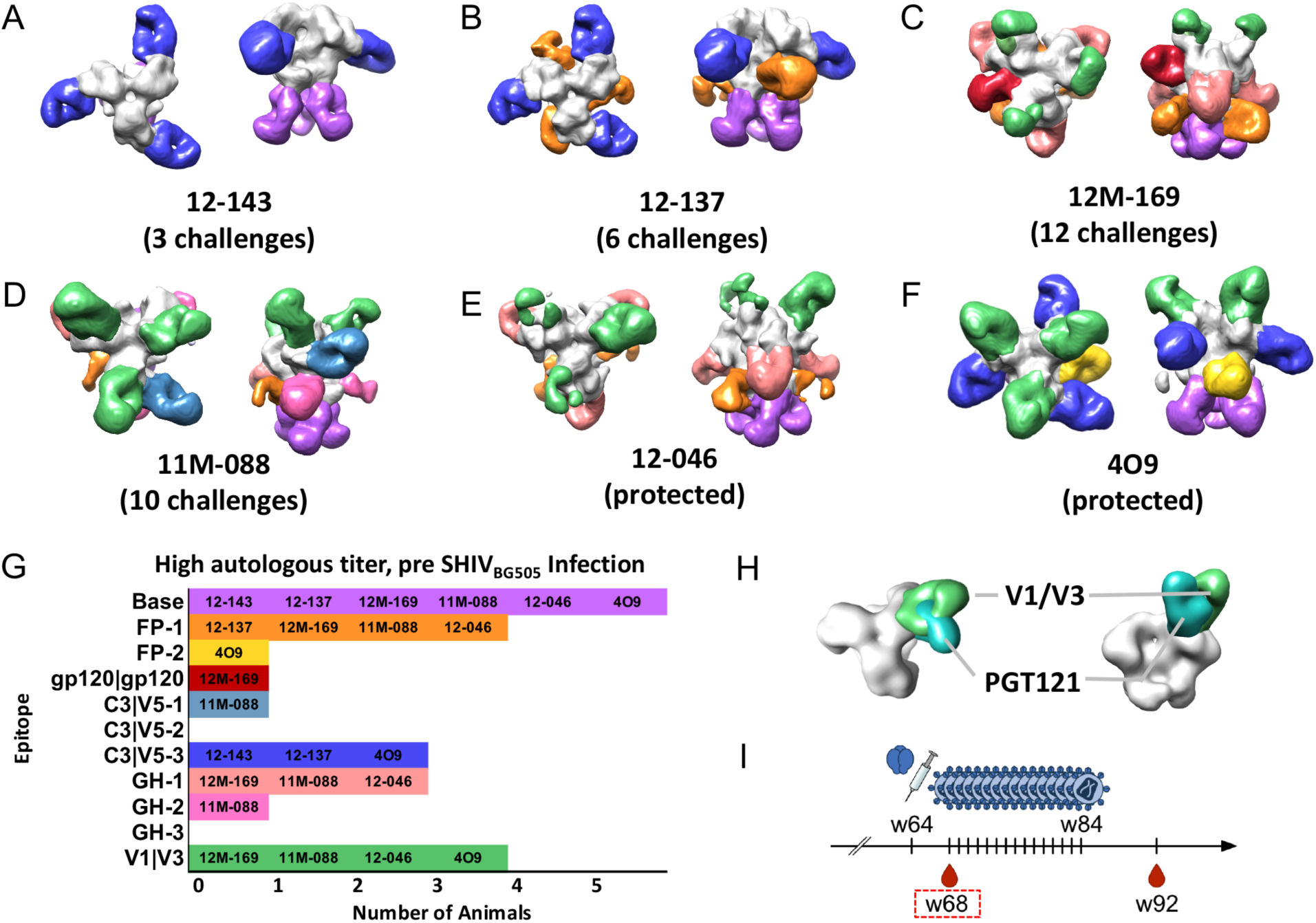
Polyclonal antibody responses of high autologous neutralizing titer macaques immediately before SHIV_BG505_ challenge. Segmented 3D reconstructions of (A) 12-143 (B) 12-137 (C) 12M-169 (D) 11M-088 I 12-046 (F) 4O9. The number in parentheses indicates the number of SHIV_BG505_ challenges it took for the animal to become infected (see Figure S3 for representative additional 3D phenotypes). (G) Bar graph summary of the epitopes targeted by high autologous neutralizing titer macaques (H) Comparison of the angle of approach between bnAb PGT121 and animal 409 V1/V3 polyclonal response. (I) The red box on timeline indicates the week from which sera were used for EM imaging. Week 68 is four weeks after the last boost.

One striking difference between the low-titer and high-titer immunized animals was the appearance of antibodies binding to the V1/V3 region near the apex of the trimer in four out of six high-titer animals compared to none of the low-titer animals. At low resolution these responses resembled known bnAbs such as PGT121 that binds to the conserved GDIR co-receptor site and the N332 glycan (Figure 3H). Another notable phenotype for the high-titer animals was a different angle of approach by which Fabs bound to the C3/V5-3 epitope: three of six animals exhibited this phenotype, with the low-titer macaques favoring the GH-1 in lieu of this putative high-titer correlate (compare Figure 2 with Figure 3A, 3B, and 3F). This observation was consistent in additional unchallenged low and high-titer animals from the immunization study (Figure S3D) (Cirelli et al., 2019; Pauthner et al., 2017). Overall, high-titer group displayed an increased diversity in the targeted epitopes, especially with the appearance of V1/V3 binding antibodies, and an associated increase in trimer occupancy (Figure 3), with apparently unique angles of approach as well, factors that may have contributed to these animals’ resistance to infection.

### Evolution of Humoral Immune Responses

Taking advantage of the intensive schedule of blood draws throughout the immunization experiment, we applied EMPEM across multiple time-points to study the evolution of antibody responses; we performed EMPEM analysis on available samples from high-titer animals after each booster immunization in an attempt to discern hierarchical signatures of developing protective antibody responses. The week 10 neutralization titers were 1:48, 1:83, and 1:404 for animals 12-137, 11M088, and 4O9, respectively (Pauthner et al., 2017). Within this subset of animals there is no clear phenotype that correlated with future ability to protect from an autologous virus challenge. For example, both 12-137 and 4O9 displayed similar epitope recognition, namely C3/V5-3 and FP but 12-137 was infected after six intra-rectal challenges while 4O9 remained uninfected after all twelve challenges. Serum samples from weeks 26-28 showed an increase in the diversity of epitopes recognized for animals 11M088 and 4O9, which was not the case for 12-137. In fact, 12-137 and another member of the high-titer group that showed only marginal protection, 12-143, did not diversify the epitopes targeted after the first boost (see also Figure 3A).

Notably, fully protected animal 4O9 mounted V1/V3-specific antibodies as early as week 26 (post-boost 2), and these were not found in 11M088, even after extensive 2D and 3D classification (Figure S4). Interestingly, we did not detect V1/V3-specific antibodies in 12-046, the other fully protected animal, at this stage, though this may have been a consequence of sample limitation and thus significantly lower Fab concentration during complex formation (see Method Details and Figure S4D). However, the weeks 66 and 68 (post-boost 3) 3D reconstructions demonstrate that both 11M088 and 12-046 ultimately generated diverse antibody responses, with V1/V3 loop now present in these animals’ polyclonal antibody responses and neutralization titers increased relative to week 26 (Figures 3E and 4B, (Pauthner et al., 2017, 2018)).

**Figure 4.**
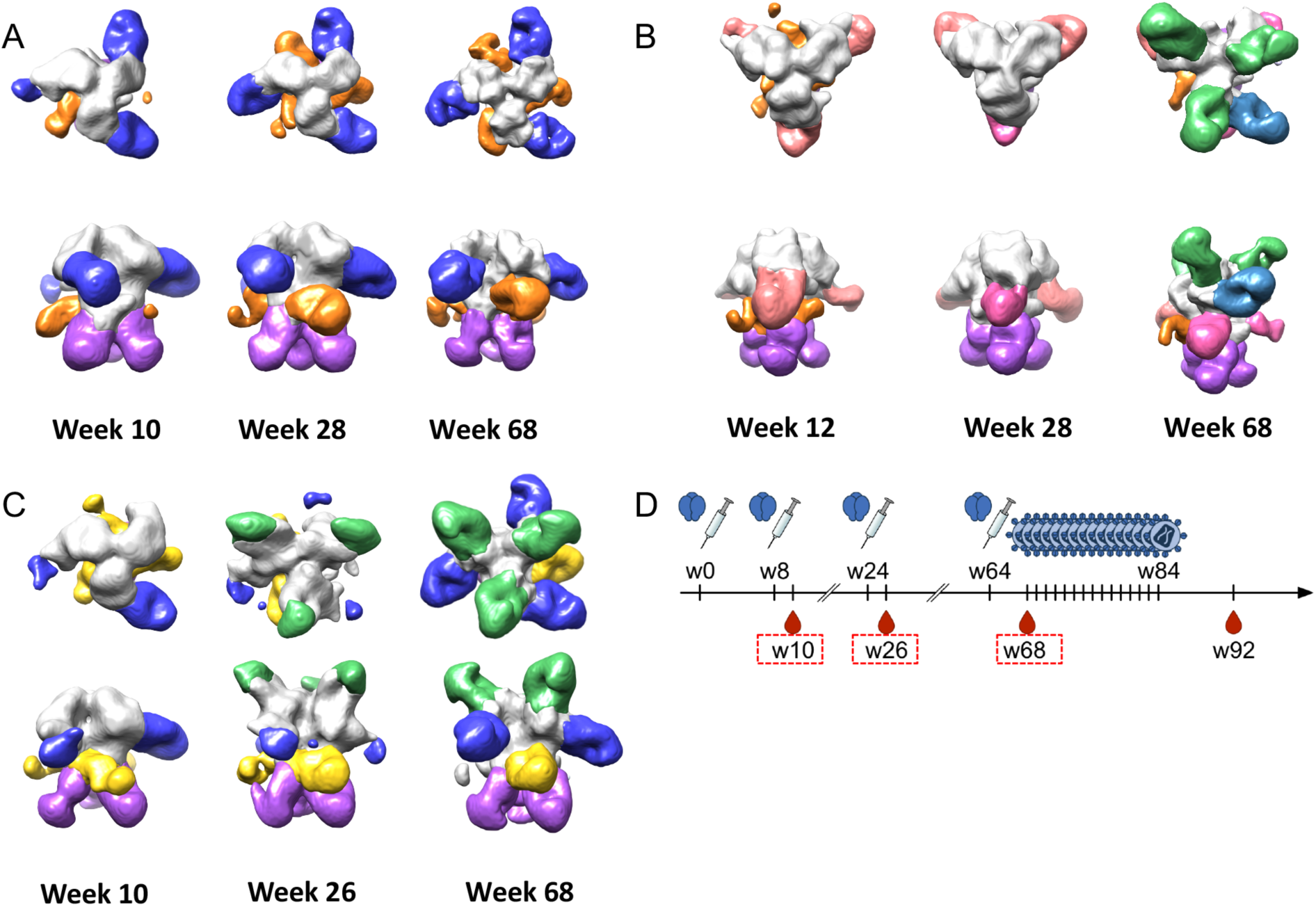
Polyclonal antibody responses of high autologous neutralizing titer macaques showing the evolution of polyclonal immune responses among animals that exhibited various degrees of protection to SHIV_BG505_ challenge. Segmented 3D reconstructions of (A) 12-137, infected after the 6^th^ SHIV_BG505_ challenge (B) 11M-088, infected after the 10^th^ SHIV_BG505_ challenge and (C) 4O9, fully protected. (D) The timeline indicates the weeks from which sera were used for EM imaging.

Thus, based on this EMPEM analyses, the ability to diversify the repertoire of epitopes recognized following continued booster immunizations generally coincided with higher neutralization titers, and protection from infection.

### Polyclonal Antibody Imaging Suggests Responses to the Fusion Peptide are Responsible for Neutralization Breadth

Pauthner et al. reported sporadic neutralization breadth in a subset of the animals’ week 26 immunization sera using the 12-virus global pseudovirus panel, with the 398F1 virus strain being the most sensitive to neutralization (deCamp et al., 2014; Pauthner et al., 2017). Notably, this virus, like BG505, also exhibits a glycan hole around position 289, which was shown to be very immunogenic, and which was recognized by antibodies observed in 12-046 (Figure 3E), among others. To determine the cross-reactive specificity(ies) we prepared complexes of 398F1 SOSIP.664 envelope trimers with week 25 or 26/28 Fabs from animals 12-137, 12-046 and 12-084 (this animal exhibited exceptionally high titers but was not among the challenged group (Pauthner et al., 2017)), and subjected them to EMPEM analysis (Figure 5A). Complexes with virus 25710 Env trimers and animal 12-046 week 26 serum were also examined (Figure S5B). These macaques had 398F1 neutralization titers of 1:32,620, 1:256, and 1:399, respectively, with 12-046 titers against 25710 at a more modest 1:160. Excluding antibodies to the trimer base, we only observed a single epitope specificity binding to heterologous trimers, namely the FP region, being contacted at two slightly different angles of approach (Figure 5A and 5B and Figure S5B). These phenotypes are reminiscent of bnAbs PGT151, VRC34, and ACS202, all of which include the FP as part of their epitope (Figure 5C, (Kong et al., 2016; Lee et al., 2016; Yuan et al., 2019)). Interestingly, the FP sequence is identical between BG505, 398F1, and 25710 (Figure S5A). Thus, based on these EMPEM analyses of a limited subset of heterologous viruses that were sensitive to neutralization by immunized macaque polyclonal sera, it appears that antibodies to the FP may be one of the main sites responsible for the breadth observed in several animals by Pauthner et al. (Pauthner et al., 2017).

**Figure 5.**
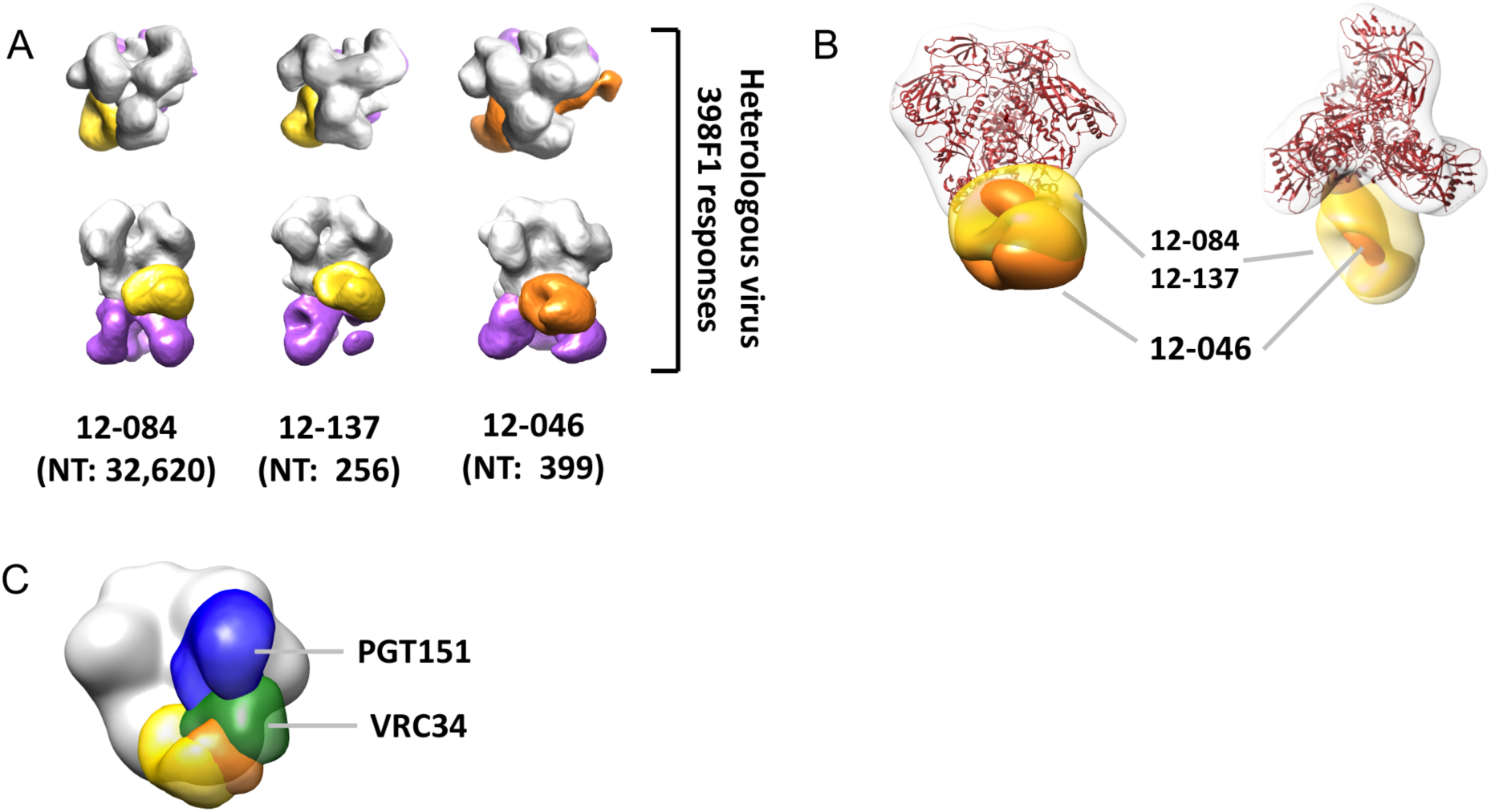
Polyclonal sera cross-reactivity mapping. (A) SOSIP version of virus 398F1 Env trimer complexed with high titer animals 12-084, 12-137, and 12-046. (B) Summary of epitopes targeted on 398F1 by high-titer animals NT = neutralization titer. (C) Comparison of polyclonal FP approach angles with bnAbs PGT151 and VRC34.

### 3D CryoEM Reconstruction Reveals Structural Details of the Macaque “V1/V3” and “C3/V5” targeting antibodies

Given the similarity of the apical antibodies that we detected in 4/6 high-titer animals to N332 bnAbs we attempted further characterization of this specificity by conducting high resolution cryoEM analysis. We used week 26 serum from the fully protected animal 4O9 (Figures 3F and 4C) to make complexes with BG505 SOSIPv5.2 to assess the structural details of the epitope recognition in high-titer animals (summarized in Figure 6). We collected 3,347 micrographs resulting in a total of 666,007 putative single particle complexes. Subsequent 2D and 3D classification using Cryosparc2 resulted in high resolution refinement of 125,800 relatively uniform complexes that produced a ∼3.9 Å global resolution 3D reconstruction (Figure 7). Notably, relative to our more typical cryoEM sample preparation conditions using mAbs, we prepared the polyclonal cryoEM sample using slightly modified conditions to maximize the likelihood of observing as many of the epitopes as possible (see Discussion and Method Details). Consequently, by attempting to maximize the Fab:trimer ratio, we generated complexes in which the Env trimers were saturated with Fabs, and consequently, we were not able to classify out trimers occupied by less than the 9+ Fabs depicted in Figure 7.

**Figure 6.**
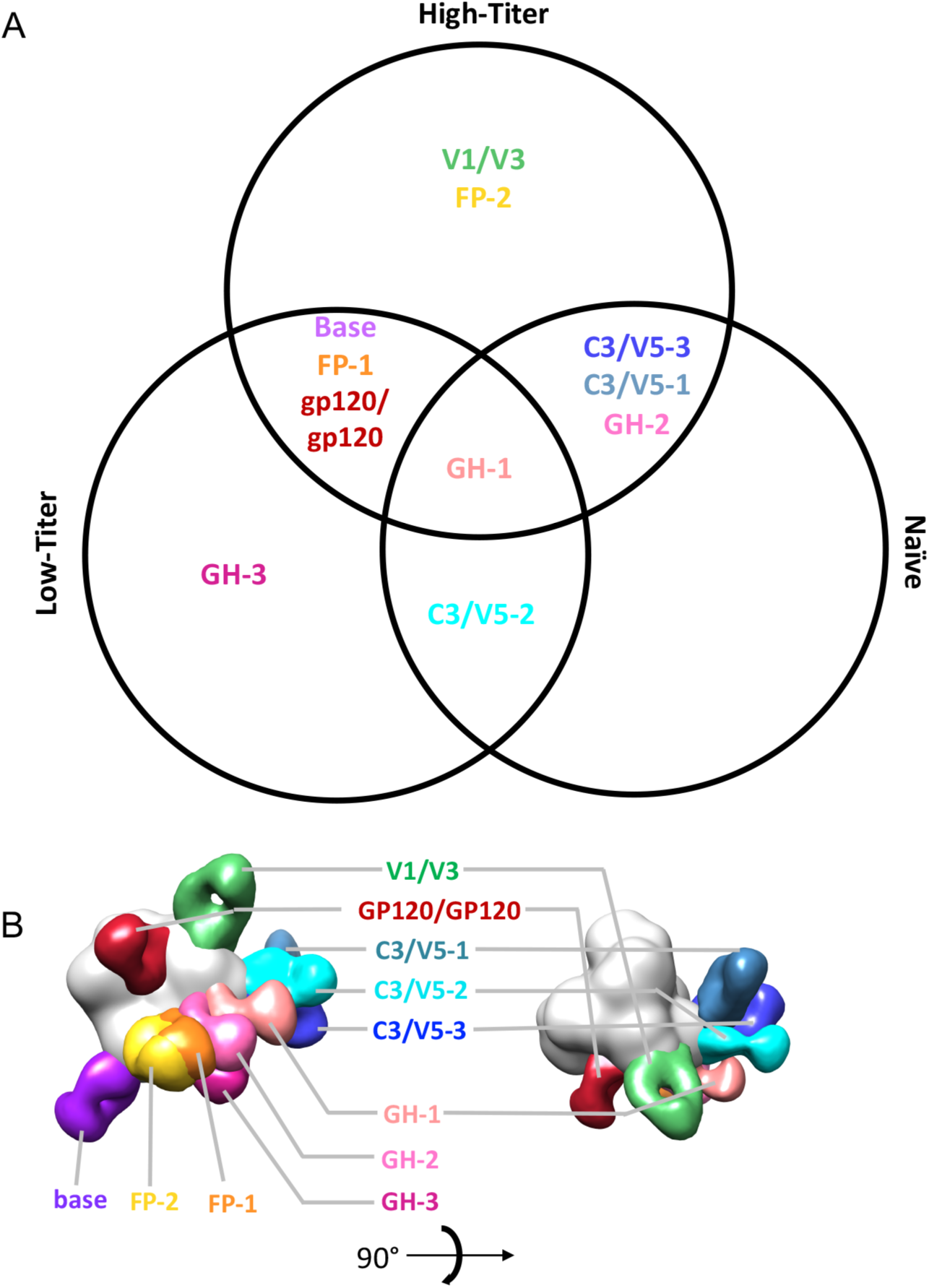
Summary of epitopes targeted by NHPs. (A) Venn diagram showing shared phenotypes among pre-challenge low-and-high-titer animals, and non-immunized infected control animals. (B) Combined global epitope summary for low, high-titer, and infected unimmunized control animals.

**Figure 7.**
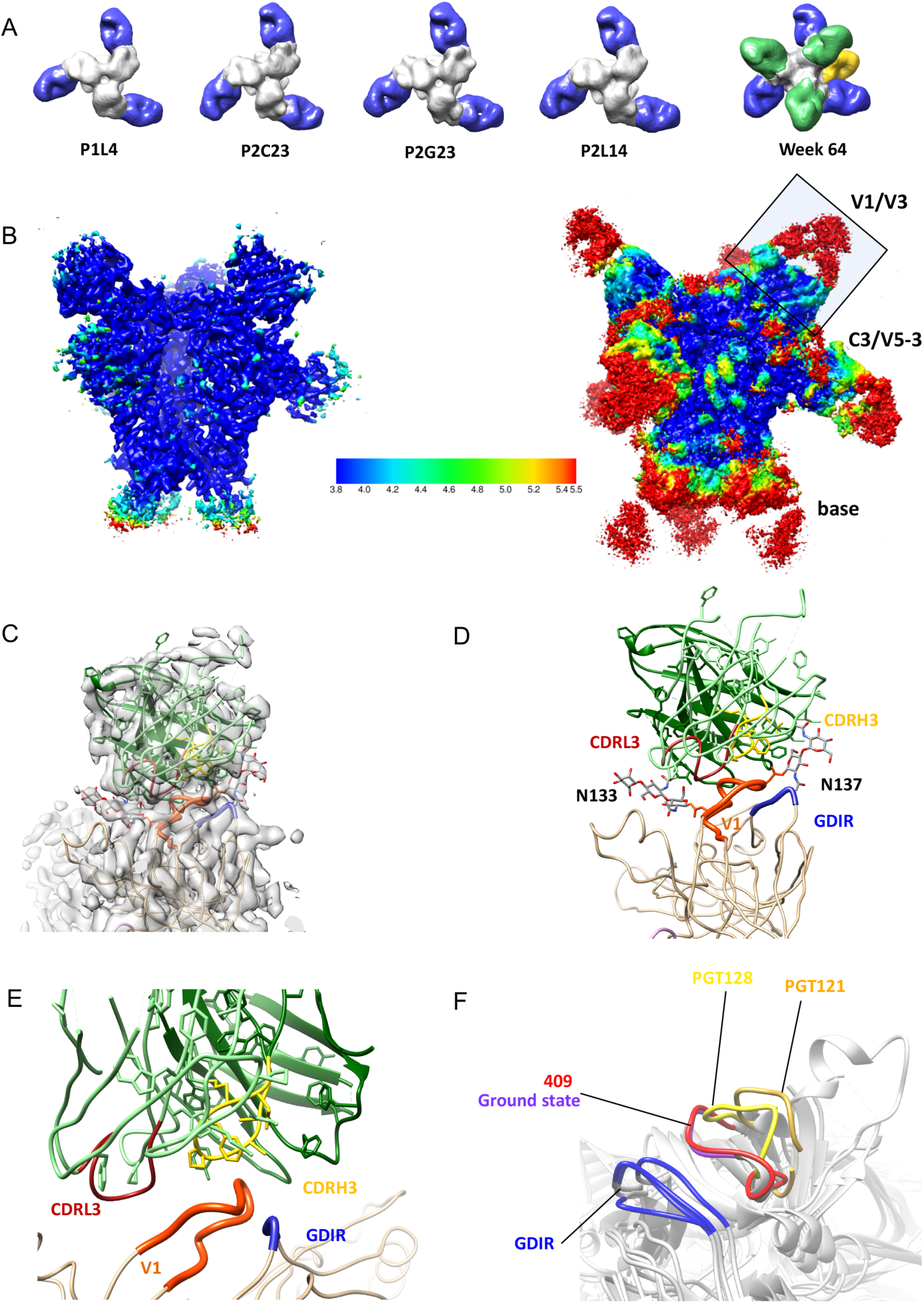
Epitopes targeted by fully protected animal 409. (A) Negative-stain reconstructions of monoclonal antibodies P1L4, P2C23, P2G23, and P2L14 isolated from animal 409. All antibodies (shown in blue) target the C3/V5 epitope similar to what we observe in the week 64 polyclonal response. (B) Low (left) and high (right) contour cryoEM resolution maps showing that the epitope-paratope regions of the V1 interactions are resolved at < 4.0 Å, with the C3/V5 and base responses showing lower resolutions. (C) and (D) Map and model of V1/V3 antibody with docked Fab demonstrating how the antibody inserts between glycans at positions 133 and 137. (E) Close-up view of V1 interaction showing proximity to and potential blocking of GDIR interaction. (F) Overlay of BG505 SOSIP.664 GDIR-V1 loop configurations of ground state (in the absence of an N332 bnAb) (PDBID: 5V8M) and with N332 glycan supersite bnAbs PGT128 and PGT121 (PDBIDs: 5ACO and 5CEZ, respectively) bound. The V1 loop in the animal 409 polyclonal complex is in the ground state, which likely precludes access to the underlying GDIR motif and therefore unlikely to develop breadth.

In the global 3D reconstruction, we observed clear density for Fabs corresponding to the trimer base, V1/V3, and C3/V5-3 epitopes. The density for the trimer base Fabs was diffuse poorly resolved, suggesting a heterogenous mixture of base antibodies were being averaged together. Conversely, the V1/V3, and to a lesser extent the C3/V5-3 Fab densities were remarkably well-resolved (Figure 7B). These data enabled more in-depth analyses of the molecular interactions of these two paratope-epitope interfaces. The V1/V3 Fabs were so well resolved that they likely corresponded to a single clonal lineage and enabled us to determine the CDR loop lengths to be ∼10, ∼14, and ∼5 residues long for CDRH3, CDRH2, and CDRH1, respectively (Kunik et al., 2012). Further, the less variable regions of the Fab were resolved to sufficient resolution to confidently build side chains.

The primary interaction between the V1/V3 Fab and the apex of the trimer is between the CDRH3 of the Fab, which navigates around glycans at positions N133 and N137 to make peptide contacts at residues N137 and I138, T139 of the trimer V1 loop, with the CDRL3 making contacts with R143 within the same region (Figure 7C and 7D). We previously reported virus neutralization data that implicated an epitope that includes residues N133 and N137, which are part of the V1 loop, as part of these antibodies’ binding (Klasse et al., 2018). Further, Pauthner et al reported up to nearly 100-fold decreases in neutralization titer in this animal upon amino acid insertions in the vicinity of these residues (Pauthner et al., 2017). Finally, the CDRH1 loop makes side-chain contacts with R327 (Figure S7H), which is a residue in the GDIR motif of the co-receptor binding site.

The V1 loop of Env in the 4O9 Fab-complexed structure is in a nearly identical conformation relative to the uncomplexed BG505 SOSIP.664 trimer control, which we deem the ground state conformation that effectively masks the conserved GDIR co-receptor binding site (Sok et al., 2016). Conversely, both PGT121 and PGT128 bnAbs binding displace the V1 from this conformation for efficient recognition of N332 and the GDIR motif (Figure 7F) (Lee et al., 2015). The cryoEM map shows that while the main interaction is between the V1 loop and the CDRH3 of the Fabs, some contacts between the V3 loop and the CDRH1 and CDRH3 of the Fabs are present, with the N332 clearly pushed out of the way (Figure S7G). Given the high sequence and glycosylation variability in V1 amongst global HIV isolates, this epitope is likely strain specific, with limited potential for broadening. This substantial epitope overlap of the NHP V1 antibodies with the N332 supersite, may constitute a roadblock for the development of bnAbs to this region.

While the less well-resolved C3/V5 epitope precluded contact residue assignment, we discerned that the primary interaction is between the BG505 variable loop 5 and the light chain of the antibody, with the glycan at position 465 shifted out of the way to accommodate the contacts (Figure S7B). The CDRH3 appears to be relatively short, possibly in order to accommodate a glycan at position 355 (Figure S7B and S7C). Notably, negative stain reconstructions of monoclonal nAbs isolated from this same animal also bound to this site (Figure 7A), with the same angle of approach as in both the cryoEM and negative-stain reconstructions of weeks 10, 26, and 64 (Figures 4C and 5A). These nAbs only partially account for the serum neutralization, further emphasizing the potential role of other epitope specificities, namely the V1 response described above.

## DISCUSSION

In an attempt to map NHP antibody responses in the context of both differing neutralization titers and levels of protection from a SHIV challenge (Pauthner et al., 2017, 2018), we applied our EMPEM approach to identify and characterize vaccine-induced antibodies that correlated with serum neutralizing titers, and consequently protection (Pauthner et al., 2018). In contrast to rabbits, where the 241/289 glycan hole and trimer base are the predominant targets of the antibody response, several additional epitopes were identified to be immunogenic in NHPs, some of which are likely responsible for the bulk of the neutralization activity (Cirelli et al., 2018; Klasse et al., 2018; Pauthner et al., 2017). These epitopes include the FP, V1/V3, and a newly identified epitope at the gp120/gp120 interface.

Antibodies elicited following SHIV_BG505_ infection of unimmunized animals revealed a relatively focused antibody response to a series of epitopes stretching from the 241/289 glycan hole to the C3/V5 region, and only comprised a subset of those observed in the immunized animals. We hypothesize two potential reasons for the differences between infection and vaccination. First, the early and immunodominant presence of base binding antibodies in the vaccine group may assist antigen recruitment to secondary lymphoid organs, and germinal centers, and/or may result in immune complexes that present the envelope trimers in a unique orientation to B-cells. Second, the persistent antigen exposure and viral escape in the SHIV infected group drives continuous evolution of focused immune responses to the two immunodominant regions on BG505 viral Env, the 241/289 glycan hole-C3/V5 super-epitope. In other words, viral infection maintains focused germinal center responses while prime-boost subunit vaccination results in more diverse epitope responses that may be influenced by the presence of circulating antibodies.

In immunized animals, high neutralization titers coincided with a more diverse epitope recognition compared to low-titer animals. Aside from one animal (12-046), this increased epitope diversity of the responses observed appeared to be associated with neutralization and protection. Neutralization was associated with epitope specificities that differed from those of the much more focused neutralizing antibody response observed in immunized rabbits (Bianchi et al., 2018). However, an increased diversity of epitopes recognized alone did not automatically result in improved protection from infection. For example, animal 12M248, which exhibited the highest diversity of epitopes in the low-titer group, did not exhibit an increased neutralization or protection (Figure 2, (Pauthner et al., 2018)), while animal 12-046 was completely protected despite a rather limited set of recognized epitopes. Also, animal 11M088 mounted antibodies to two more epitopes relative to the fully-protected animals and one more relative to 12M169, yet this animal was infected the earliest of all the animals that targeted the V1/V3 epitope (10 challenges). Interestingly, two of the four animals in the low titer group that became infected after only two challenges had more diverse responses than those that became infected after a single challenge (Figure 2E). Thus, while diversification of recognized epitopes may increase the likelihood of neutralization or protection, the overall titers of neutralizing antibodies appeared to be primarily responsible for protection.

C3/V5 was highly immunogenic, and we found a range of approach angles for antibodies binding to this epitope. Some, namely those designated C3/V5-3 antibodies, were only detected in high-titer animals (Figure 3), a finding that is consistent with a prior study (Cirelli et al., 2018). Still, without the additional presence of V1/V3 antibodies, the C3/V5-3 epitope by itself was insufficient for durable protection in the face of repetitive SHIV challenges. Indeed, viruses isolated from animal 12-137, which became infected after six challenges, displayed putative escape mutations at positions 354 and 356 (Pauthner et al., 2018). Since these residues that flank the N355 glycan, which is part of the C3/V5-3 epitope (Pauthner et al. 2018), it appears that either the escape threshold for this epitope is relatively low, or that antibodies against this epitope exert a strong selective pressure for escape. Also, given its overlapping footprint with the CD4bs, C3/V5-3 class antibodies will compete with and potentially suppress the induction of CD4bs bnAbs like VRC01.

Despite being a potential correlate of protection in this NHP challenge model our cryoEM analysis reveals that the V1/V3 class of antibodies is likely competitive and counterproductive to the induction of N332 supersite bnAbs (Figure 7E). Should the human humoral response to the trimer apex be analogous to rhesus macaques, our high-resolution structure suggests that additional engineering may be necessary to optimize the N332/GDIR epitope presentation in this immunogen.

While detection of FP-like antibodies in rabbits was rare (Bianchi et al., 2018), FP responses were readily detectable in rhesus macaques as early as week 10. We also show that FP epitope likely contributed to the observed neutralization breadth in some animals, and is therefore the most promising antibody response elicited by the BG505 trimer immunization. Further, because the FP is a well-characterized target of human bnAbs, our results suggest that macaques are a suitable preclinical model for evaluating FP-directed immunogens intended to elicit PGT151-ACS202- or VRC34-like bnAb.

With the first dose-escalation Phase I clinical trial using the BG505 SOSIP.664 gp140 trimer as the immunogen now recruiting human subjects (McElrath, Julie, Omu Anzala, 2018), the present findings are an important preclinical benchmark. We show that, similar to rabbits, the macaque preclinical model develops polyclonal responses in a tiered fashion, with antibodies against the most accessible immunogenic epitopes elicited early, followed by more diverse and individualized responses. We predict that human responses will be similar to NHPs, and our EMPEM approach will enable rapid comparison to this end. Moreover, the present structural information can be exploited to design epitope knock-out trimer probes that can be used to quantify epitope-specific B cell responses by FACS, or that can be used for the isolation of B cells subsets specific for individual epitope(s); a more detailed characterization of monoclonal antibodies will offer further lessons for immunogen redesign. Finally, our detailed structural analyses identified potential weak points of the current generation of BG505 SOSIP trimer immunogens and will help to inform the design and engineering of future iterations of this immunogen.

## ACKNOWLEDGMENTS

This work was supported by NIH grants R01 AI136621 and UM1 AI100663 (ABW and LH), Swiss National Science Foundation grants P300PB_160969 and P3P3PB_160970. C.A.C. is supported by a NIH F31 Ruth L. Kirschstein Predoctoral Award Al131873 and by the Achievement Rewards for College Scientists Foundation.

## AUTHOR CONTRIBUTIONS

(B) N. designed and performed the electron microscopy experiments and performed the analysis. M.B. prepared the complexes for nsEM. C.A.C. and L.M.S. assisted B.N. with sample preparation and provided SOSIP trimers. H.L.T. assisted with cryoEM data collection. R.N.K. assisted with model building and refinement. D.S. and F.Z. performed the B-cell sorting and provided monoclonal antibodies. D.R.B. advised on the project and the manuscript. A.B.W. and L.H. designed and supervised the project. B.N., A.B.W. and L.H. wrote the manuscript. All authors were asked to comment on manuscript.

## STAR★METHODS

### KEY RESOURCES TABLE

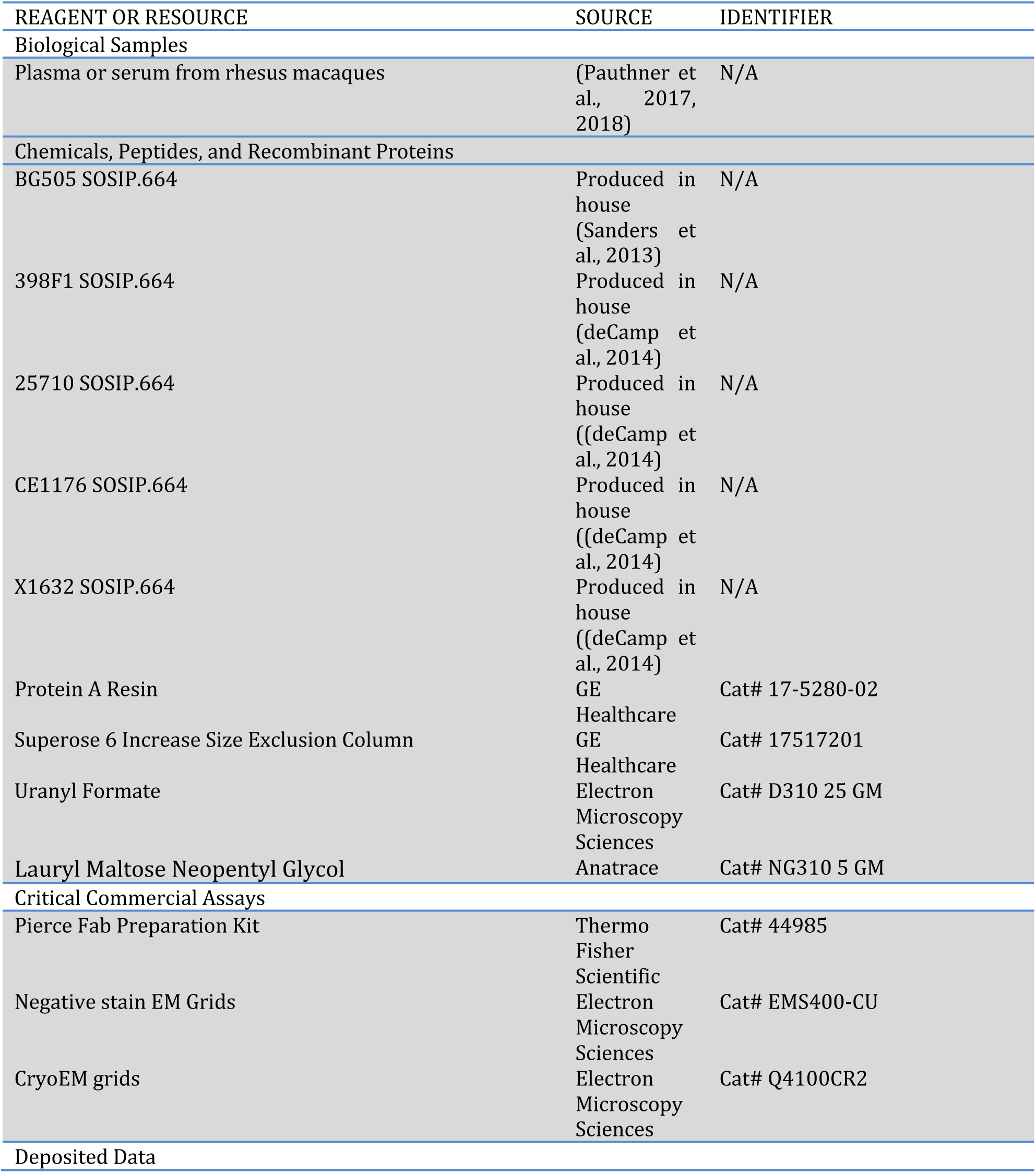
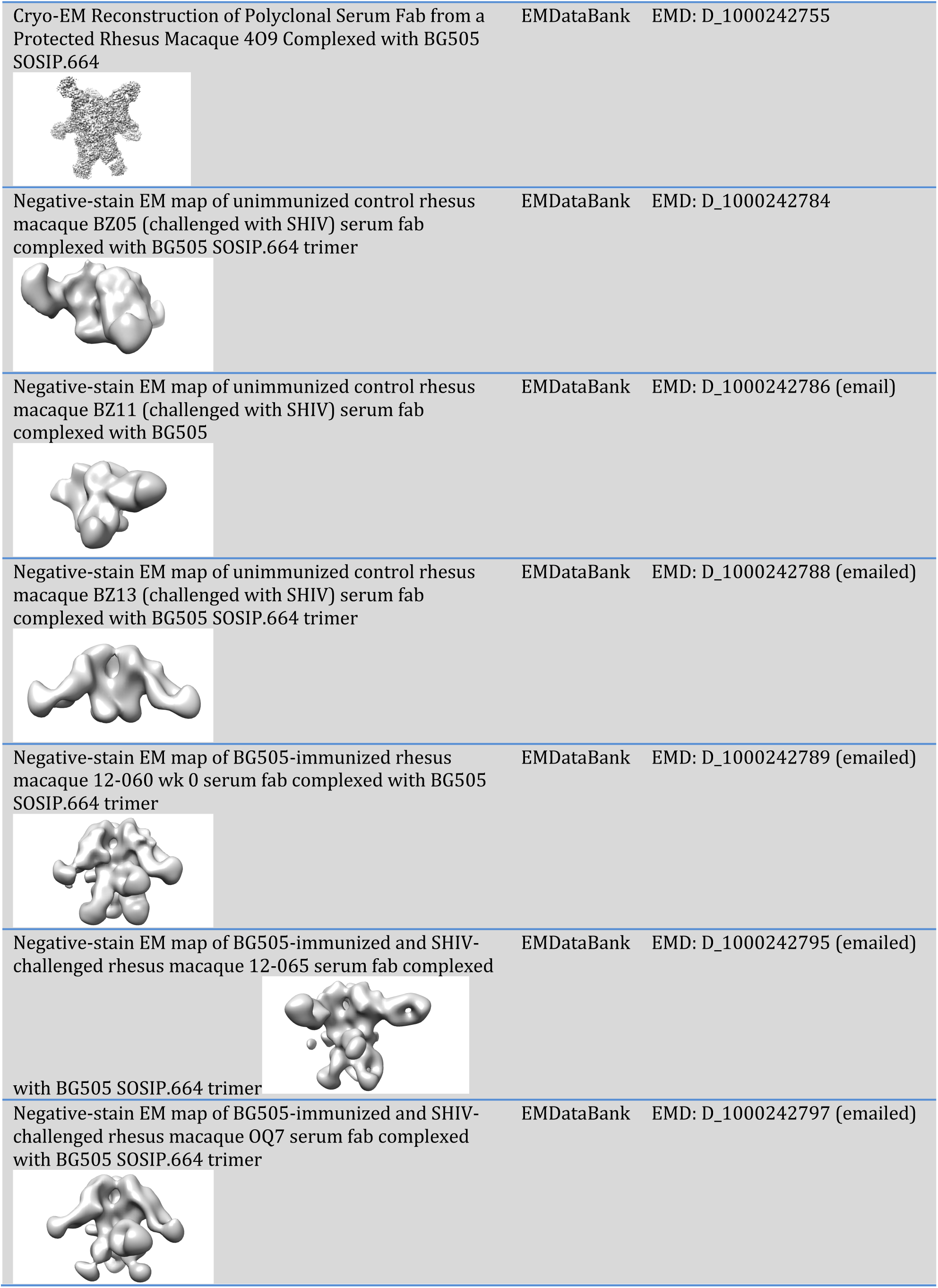
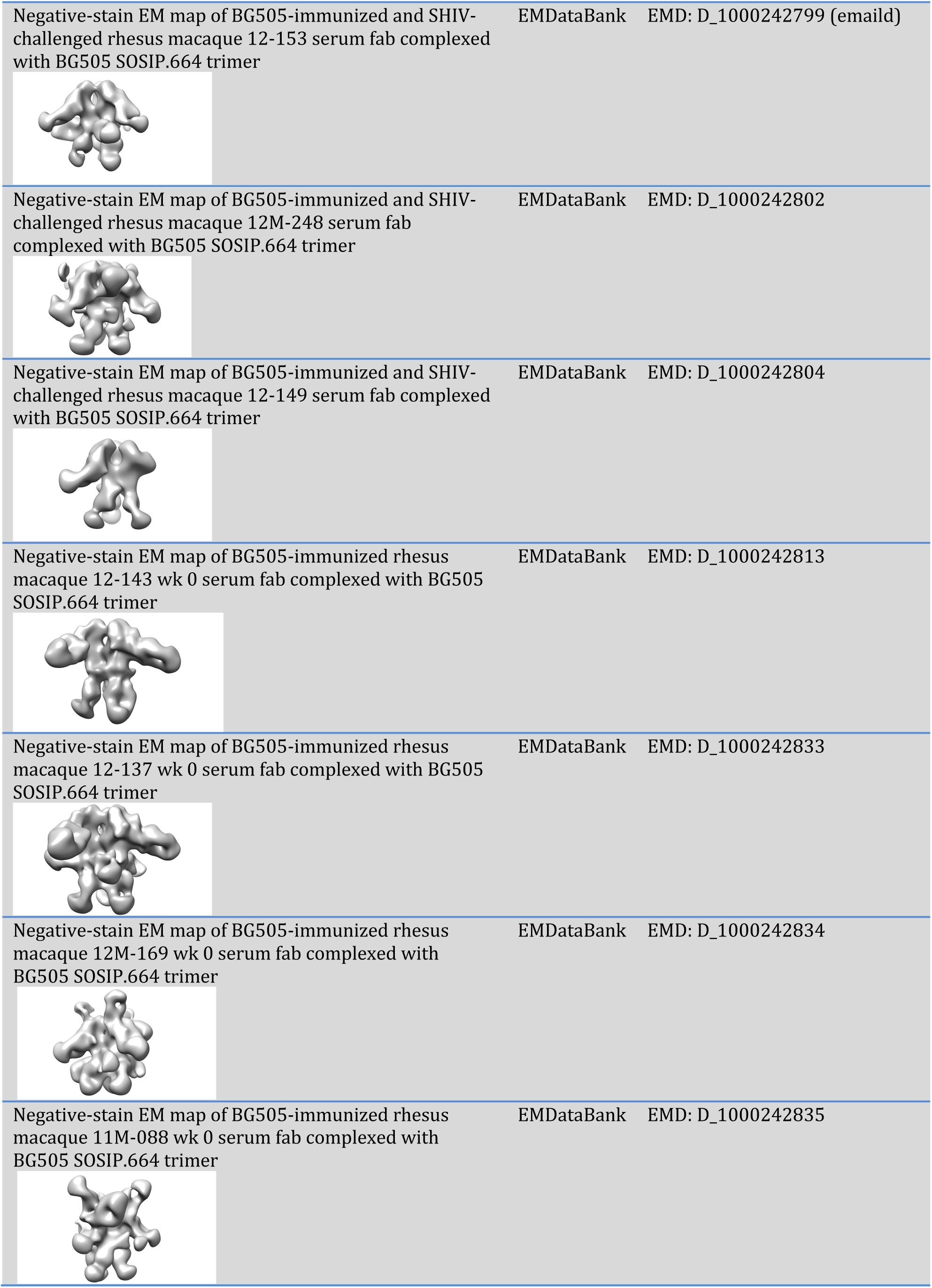
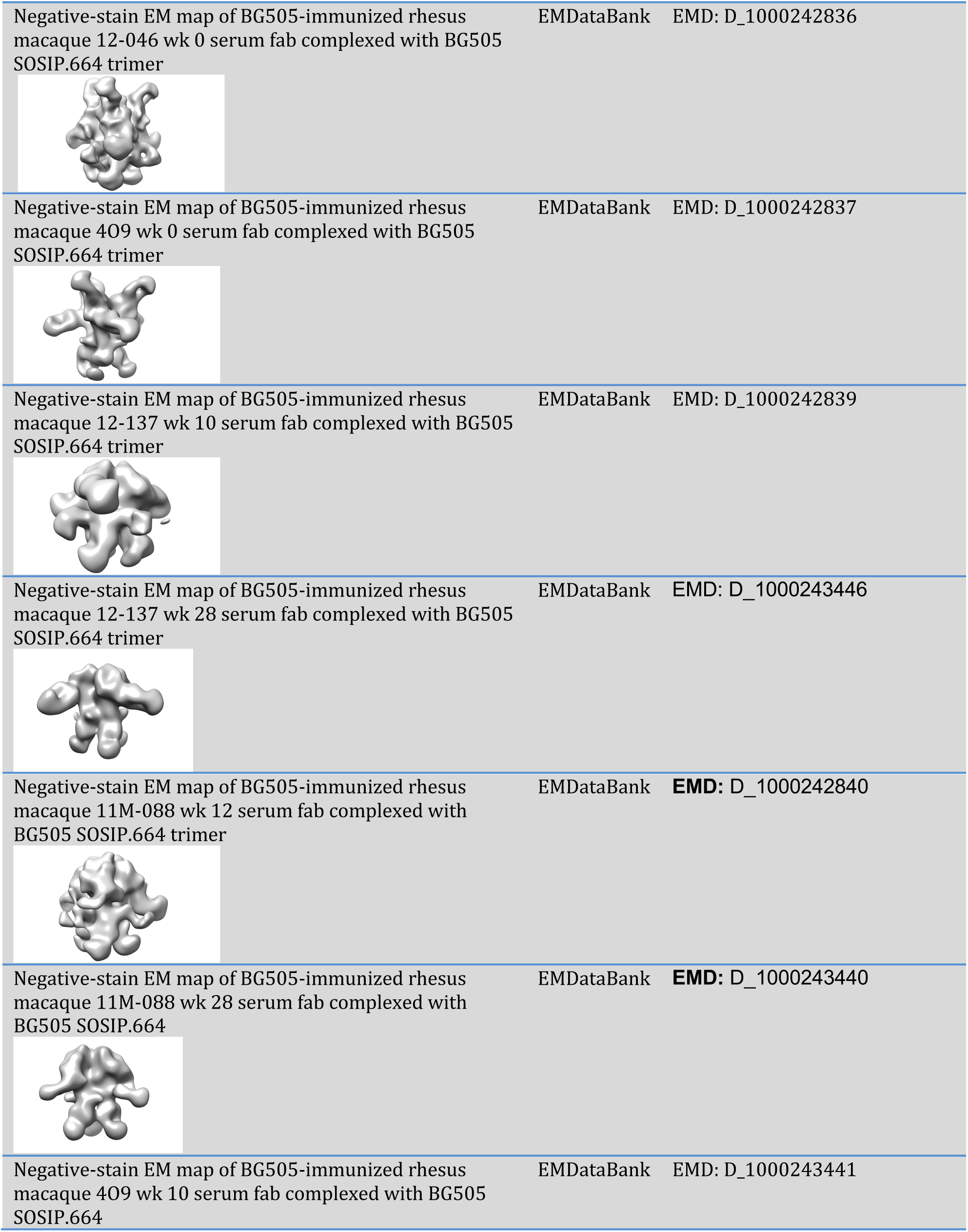
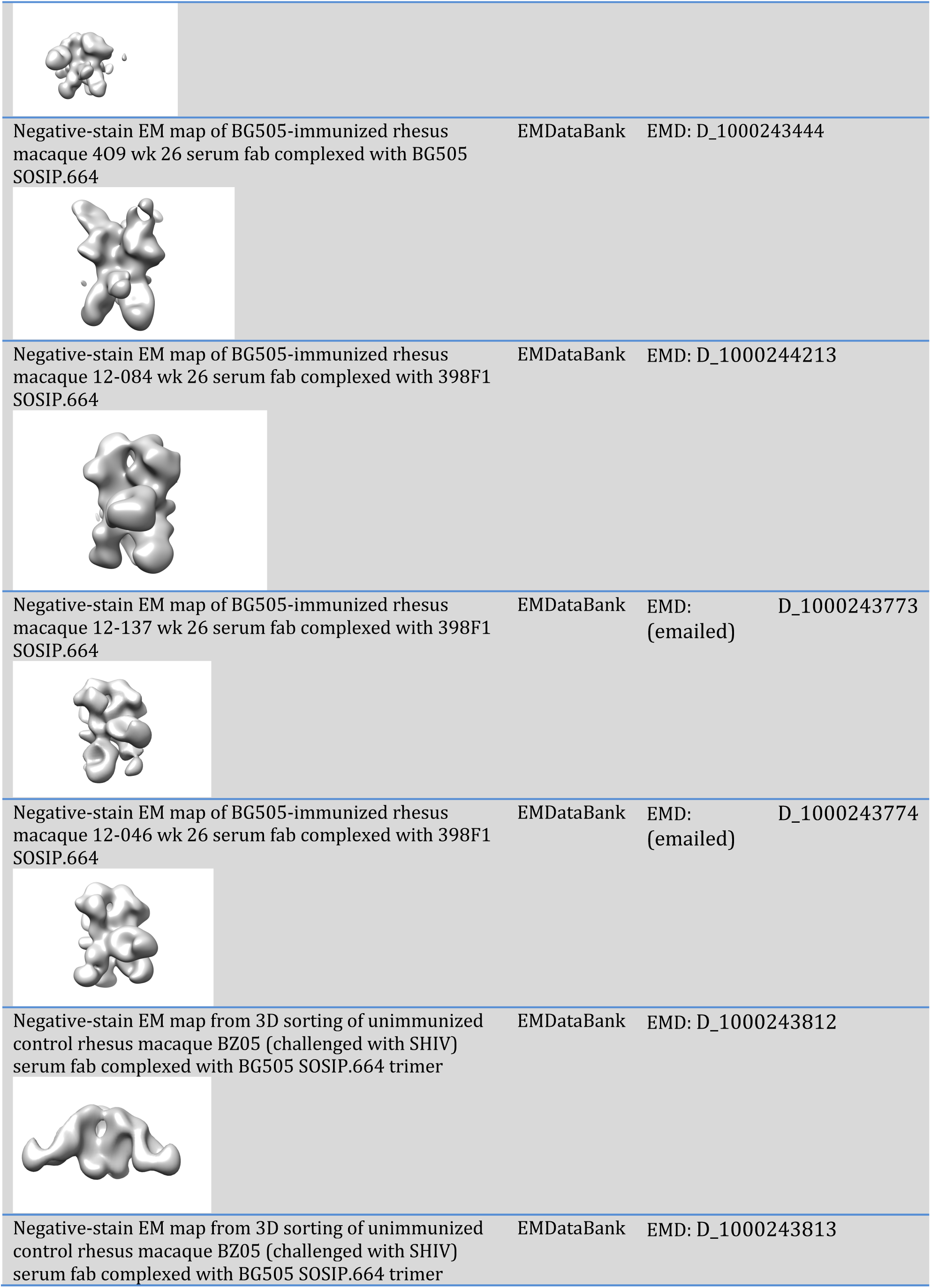
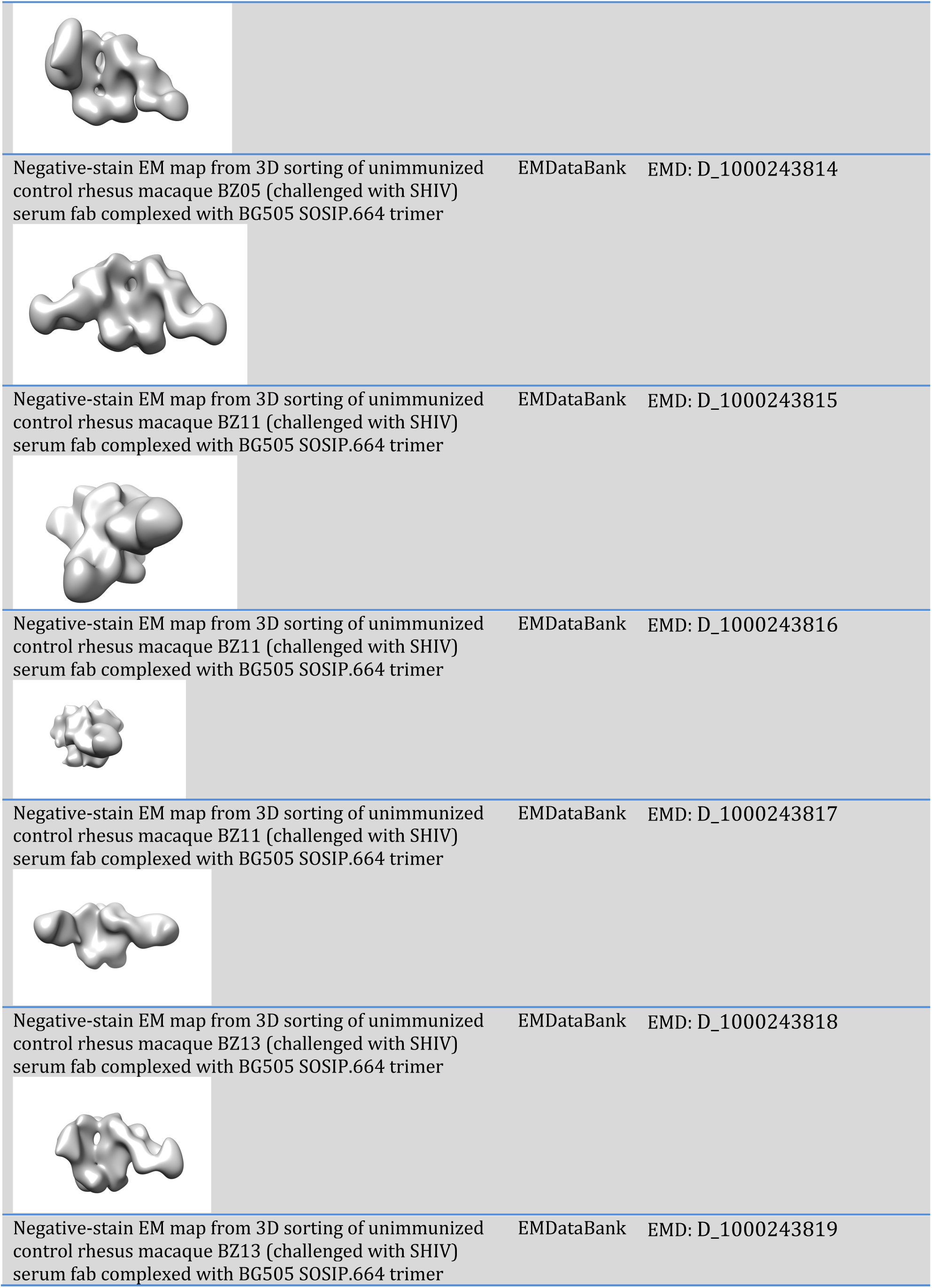
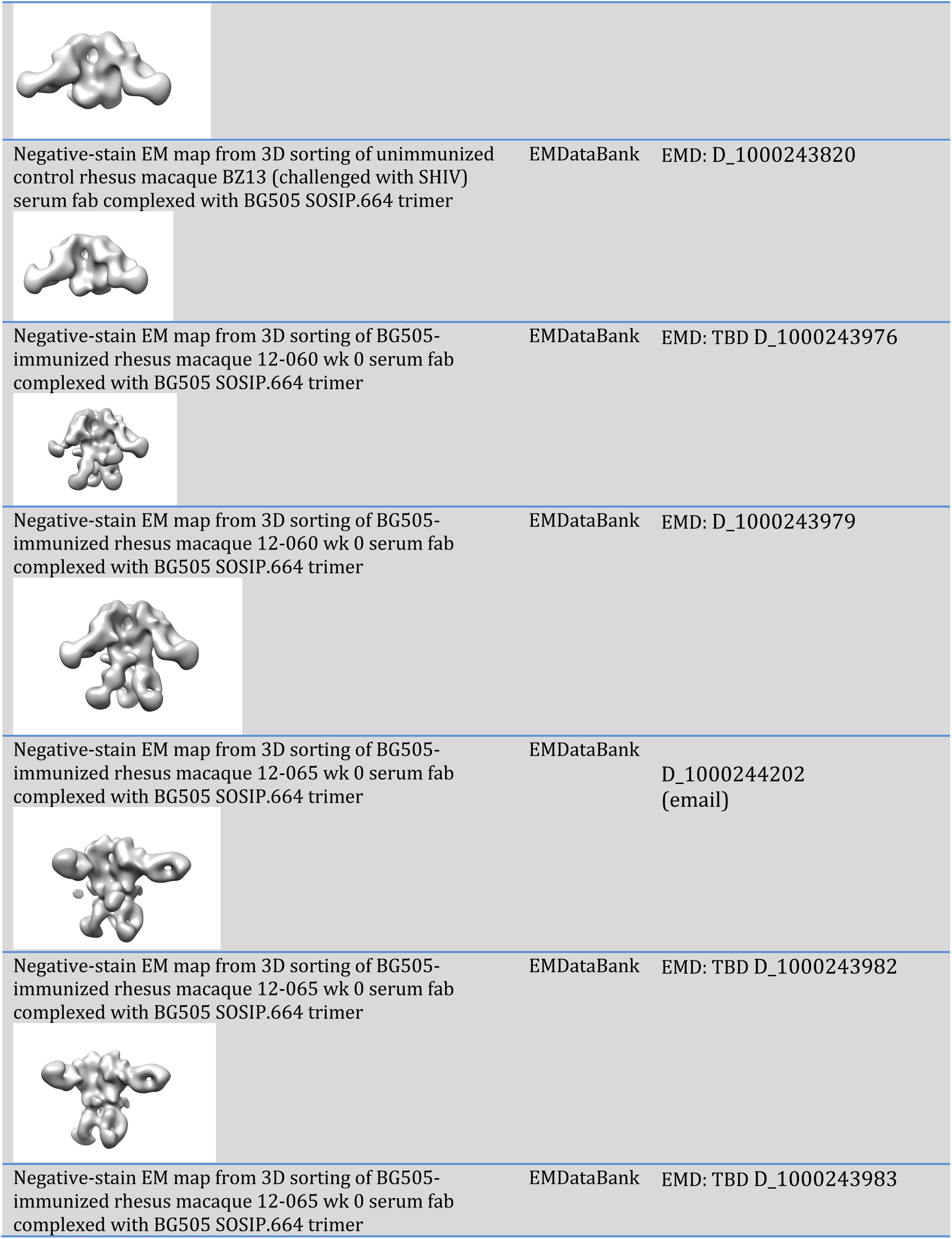
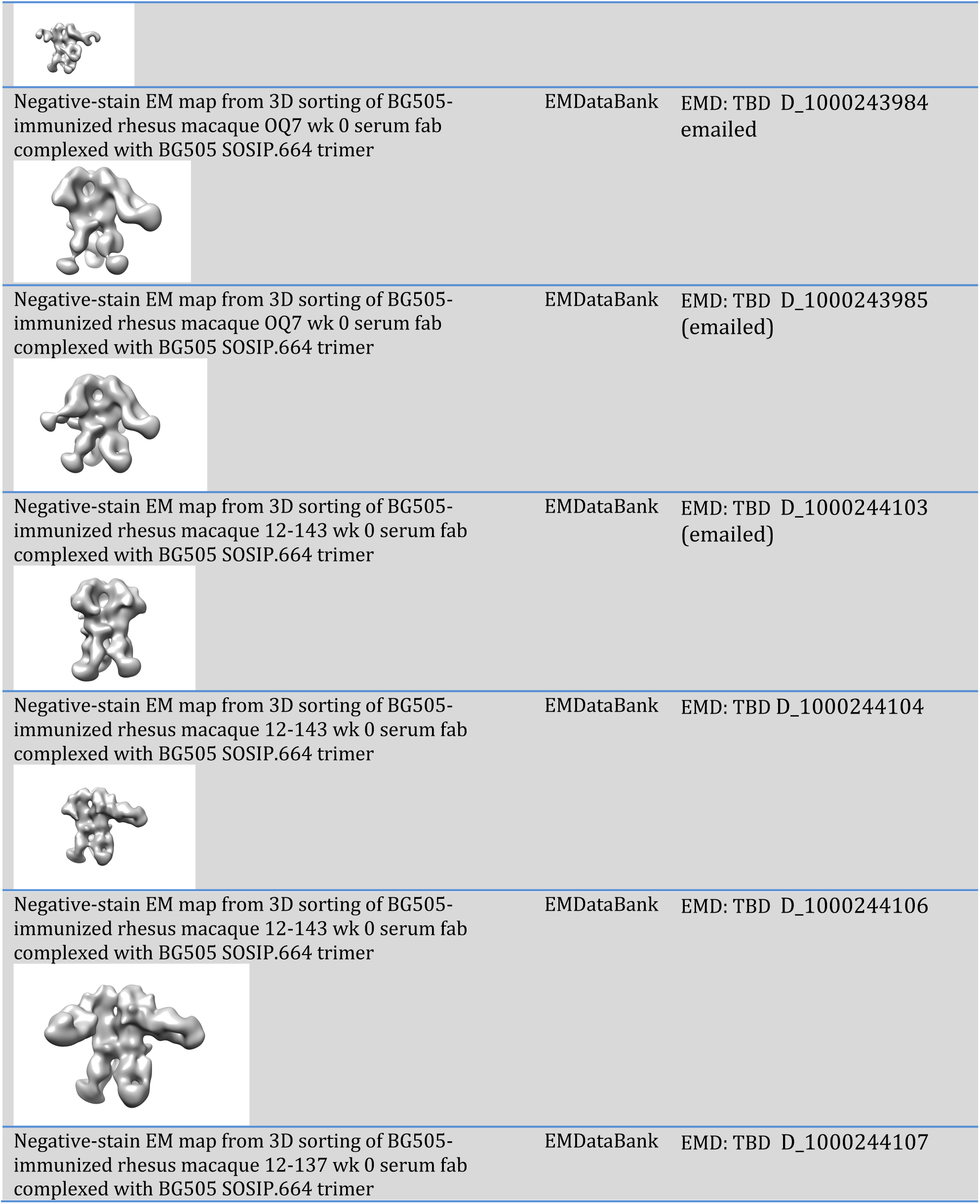
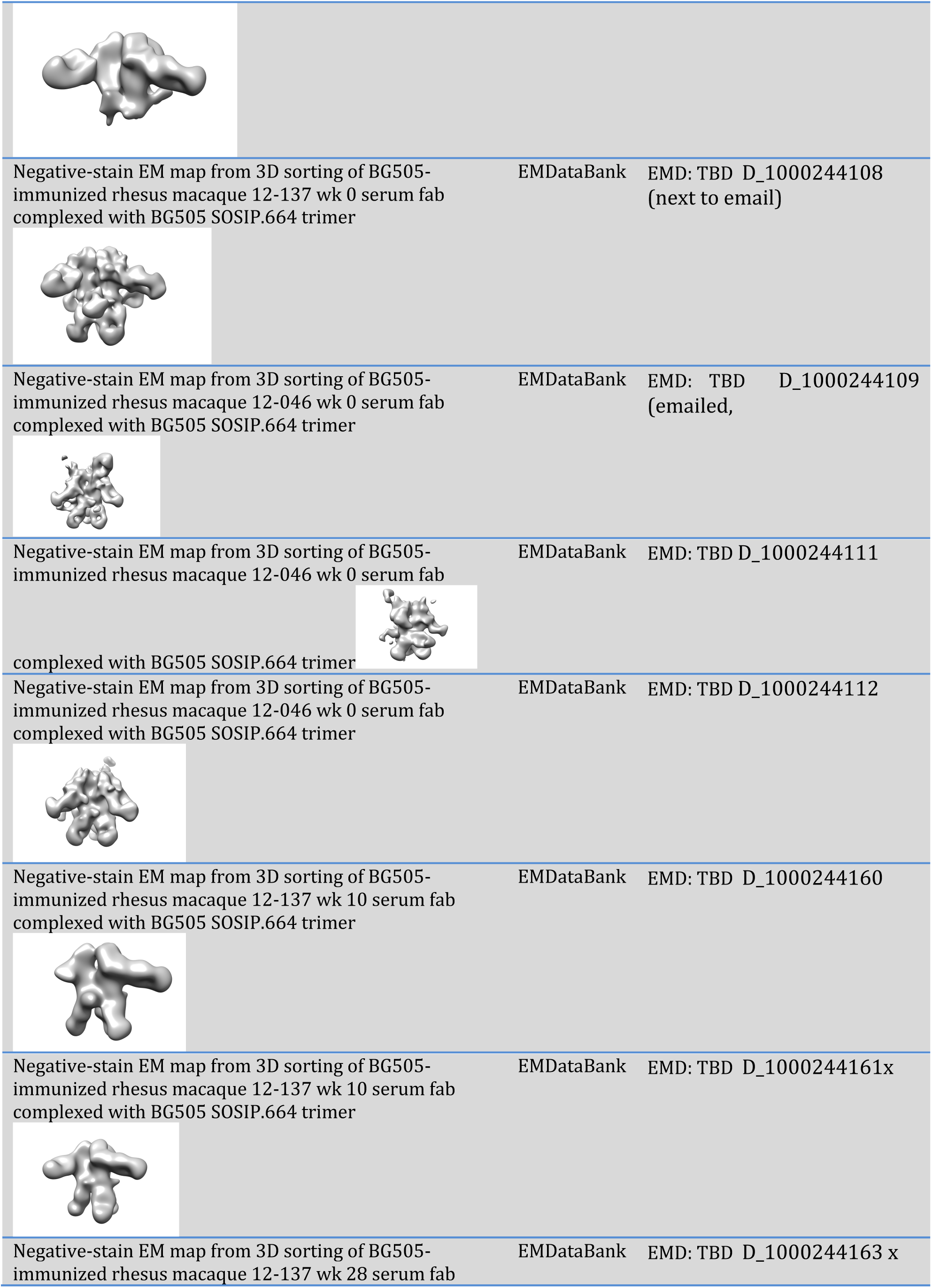
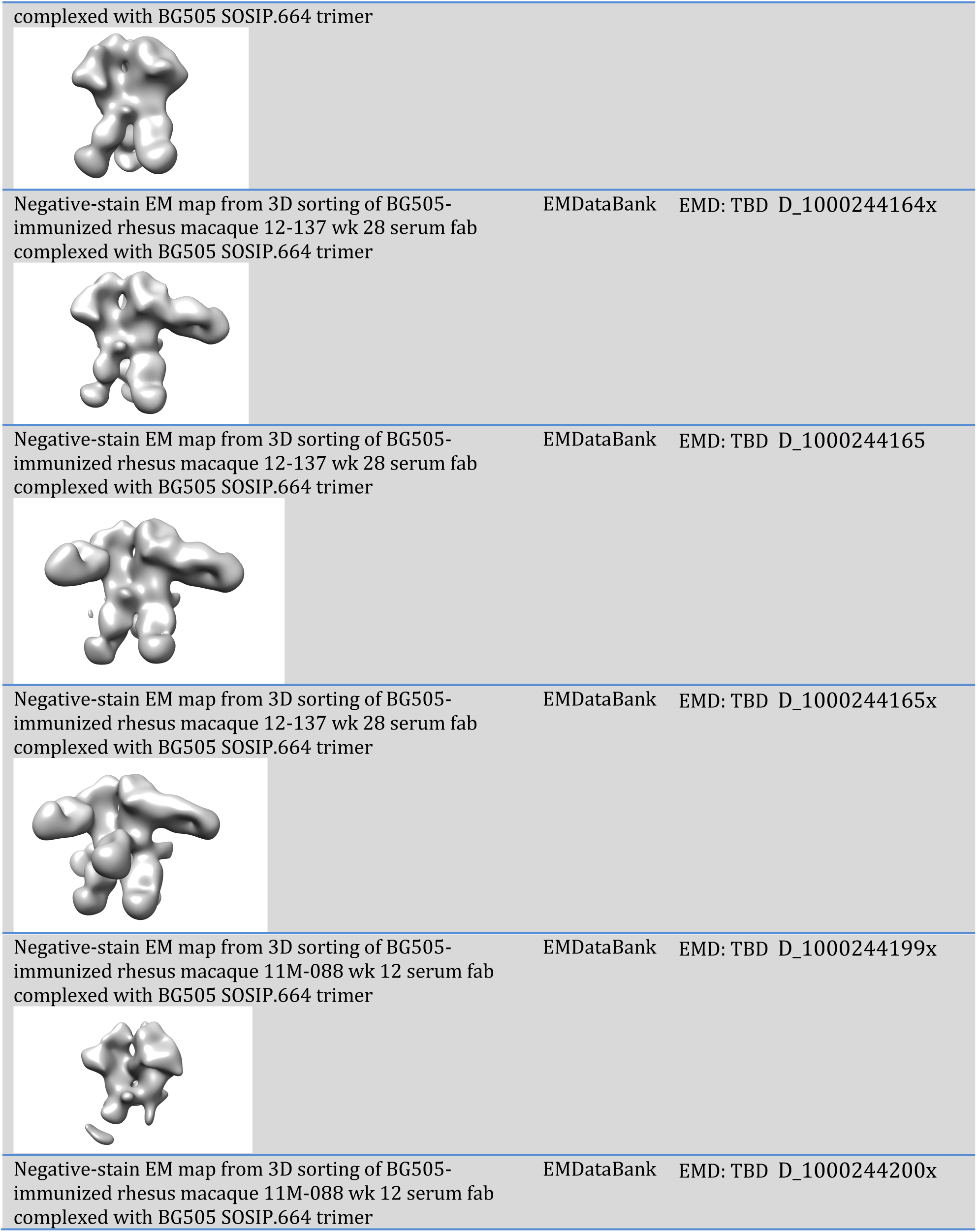
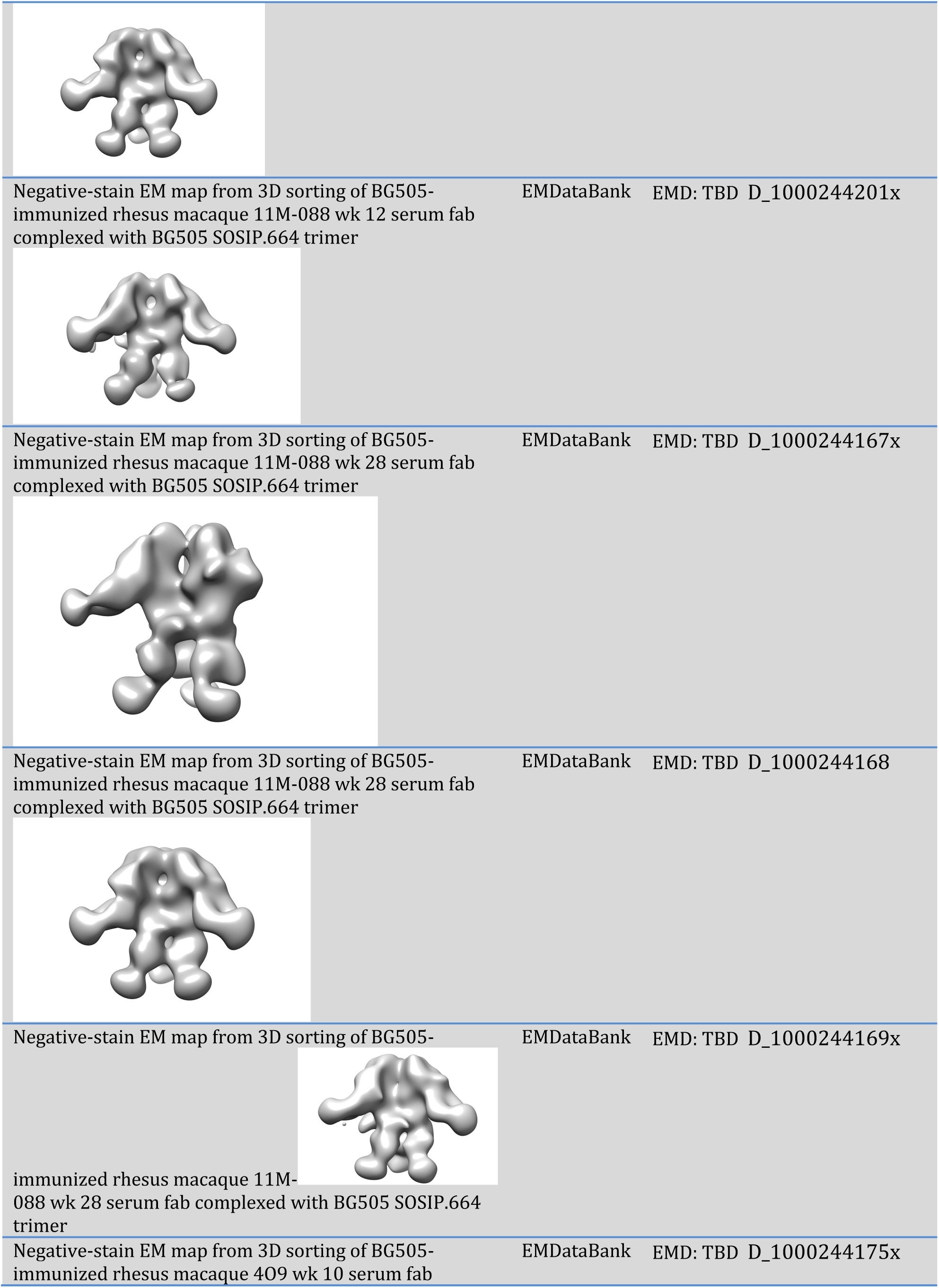
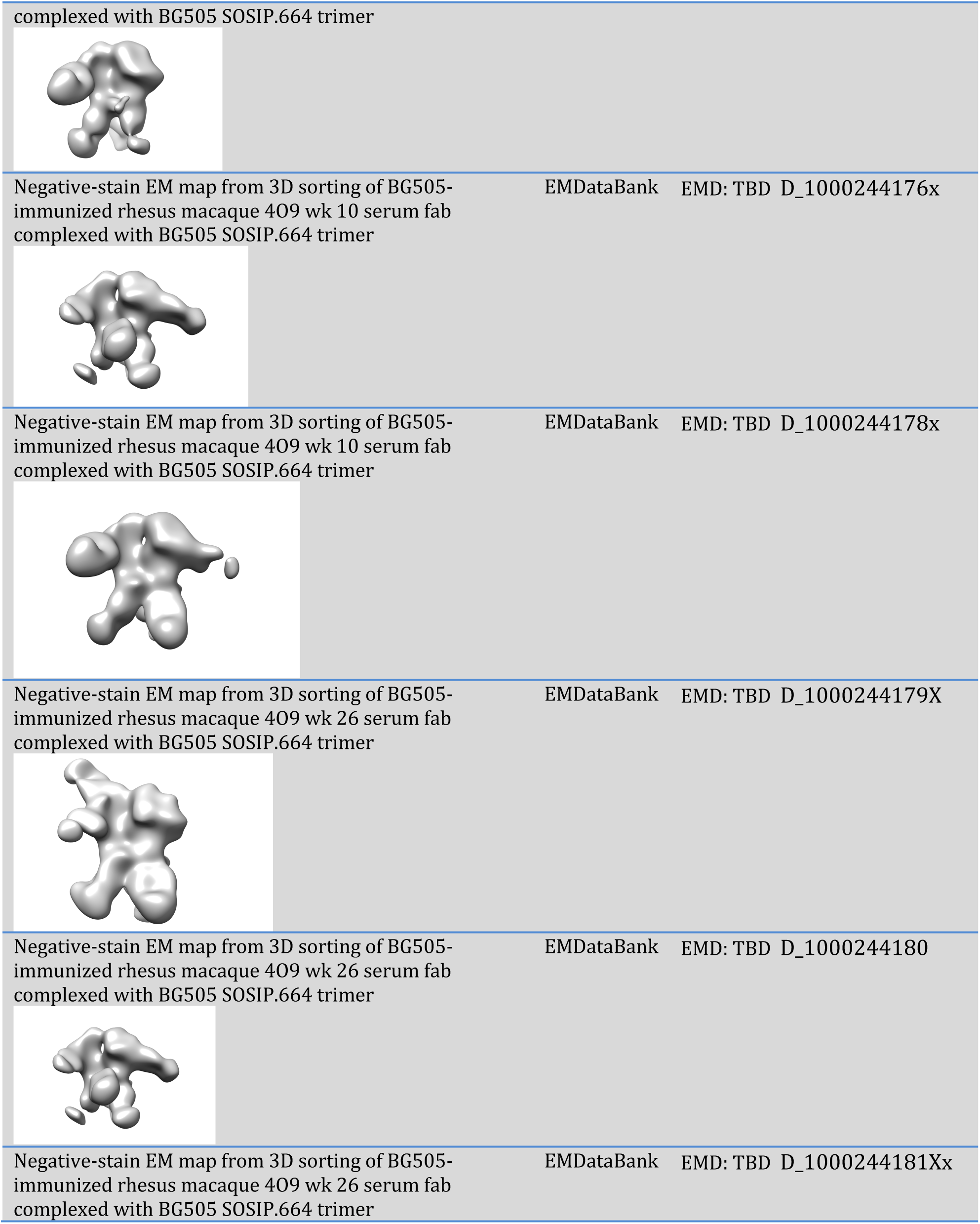
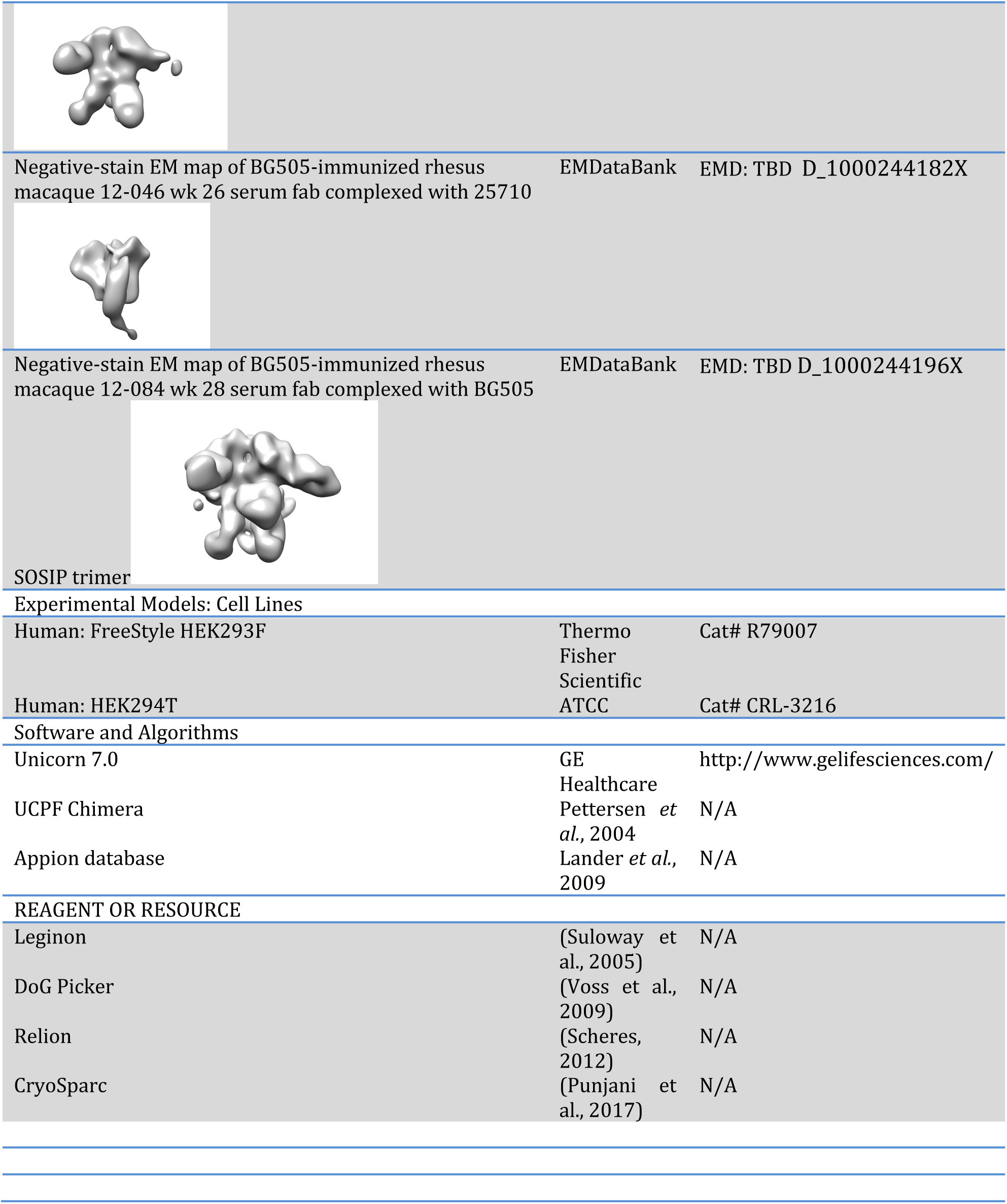

### CONTACT FOR REAGENT AND RESOUCE SHARING

Further information and requests for resources and reagents should be directed to and will be fulfilled by the Lead Contact, Andrew B. Ward

### EXPERIMENTAL MODEL AND SUBJECT DETAILS

#### Rhesus Macaques

The plasma and serum samples used in this work derive from the previously described rhesus macaque immunization and subsequent challenge experiments using a subset of the cohort (Pauthner et al., 2017, 2018). Briefly, 70 rhesus macaques received a prime and 2 booster immunizations over the course of 26 weeks. Of this immunized cohort, 6 high and 6 low neutralization titer animals were selected to enter the challenge study, with 6 non-immunized (“naïve”) animals used as controls. The Scripps Research Animal Care and Use Office and the Committee (IACUC) approved all experimental procedures involving all the animals.

### METHOD DETAILS

#### Soluble Env Protein Production

BG505 SOSIP.664 v5.2, 25710, X1632, CE1176, and 398F1 SOSIP.664 were co-transfected with furin into HEK293F cells, followed by affinity chromatography using either 2G12 or PGT135 as the ligands. Affinity eluents were then purified via size exclusion chromatography (SEC) in order to remove trimer aggregates or protomers. Full details of trimer purification can be found elsewhere (Pugach et al., 2015).

#### Plasma or Serum IgG Purification

Serum or plasma polyclonal IgGs were Protein A affinity-purified using a 1:1 ratio of resin to undiluted serum or plasma sample. After a 4-to-5-fold dilution with PBS, samples were incubated at room temperature (RT) for 5 h or overnight at 4 °C, followed by a 3-fold wash step using 10 volumes of PBS. Elution consisted of 5-10 volumes of 0.1M glycine, pH 2.5 into vessels pre-conditioned with ∼20% v/v Tris-HCL, pH 8. Eluents were buffer exchanged into TBS by centrifugation with 10 kDa cutoff membranes (EMD Millipore).

#### IgG digestion

In order to perform EM imaging, Fabs were produced by digesting the IgGs using resin-immobilized papain (50 μl settled resin/mg IgG, Thermo fisher Scientific). The digestion occurred in 20 mM sodium phosphate, 10 mM EDTA, 20 mM cysteine, pH 7.4 for 5-6 h at 37 °C. In order to remove Fc and intact IgG, the digestion mix was incubated with protein A Sepharose (GE Healthcare) for 1 h at RT. The Fab-containing supernatant was buffer exchanged into TBS by centrifugation with 10kDa cutoff membranes (EMD Millipore).

#### EM Complex Preparation

Complexes for negative-stain EM were prepared similarly to the previously described method, except ELISA EC50 was not used to guide serum Fab concentration to be used (Bianchi et al., 2018). Briefly, after buffer exchanging into TBS, up to ∼1 mg of total Fab was incubated overnight with 10 μg BG505 trimers at RT in ∼50 μl total volume. Complexes were then purified by SEC using Superose 6 Increase 10/300 column (GE Healthcare) in order to remove unbound Fab. The flow-through fractions containing the complexes were pooled and concentrated using 100 kDa cutoff centrifugal filters (EMD Millipore). The final trimer concentration was titrated to ∼0.04 mg/mL prior to application onto carbon-coated copper grids. Of note, depending on sample availability, not all complexes were prepared using identical Fab concentrations during incubation with trimers, with samples from animals where serum material was limiting resulting in lower Fab:trimer ratio. For animal 12-046, week 10 and 26 samples were insufficient to include in the time-course study as the purified total Fab concentration was severalfold lower than the rest of the animals (see table below).

| Sample ID | Fab concentrations (mg/mL) |  |
| --- | --- | --- |
|  | Week 10 | Week 26 |
| 12-137 | 14 | 35 |
| 11M088 | 29 | 33 |
| 409 | 37 | 27 |
| 12-046 | N/A | 4.8* |

#### Negative-stain EM

The SEC-purified complexes were applied to glow-discharged, carbon-coated 400-mesh copper grids, followed by pipetting 3 μl of 2% (w/v) uranyl formate stain and blotting, followed by application of another 3 μl of stain for 45–60 s, again followed by blotting. Stained grids were stored under ambient conditions until ready for imaging. Images were collected via Leginon software using a Tecnai T12 electron microscopes operated at 120 kV and 52,000x magnification (Suloway et al., 2005). In all cases, the electron dose was 25 e^−^/Å ^2^. Particles were picked from the raw images using DoG Picker and placed into stacks using Appion software (Lander et al., 2009). 2D reference-free alignment was performed using iterative MSA/MRA. The particle stacks were then converted from IMAGIC to RELION-formatted MRC stacks and subjected to RELION 2.1 2D and 3D classification (Scheres, 2012).

#### CryoEM

Animal 4O9 complex preparation for cryoEM was modified relative to the negative stain workflow in order to maximize trimer occupancy while still resulting in sufficient complex concentration for cryoEM grids. Here, ∼50 μg of trimer was incubated with >3 mg of Fab and incubated overnight as indicated above. The complex was then concentrated to 1.5 mg/mL and mixed with Lauryl Maltose Neopentyl Glycol (LMNG, Anatrace) prior to deposition onto 2/2 Quantifoil grids (EMS) that were glow discharged for 10 s directly preceding the deposition in a Vitrobot (Thermo Scientific). Once sample was deposited the grids were blotted and plunged into liquid ethane using the Vitrobot to immobilize the particles in vitreous ice. Using Leginon image acquisition software we collected 3,347 micrographs at a nominal magnification of 29,000x with a Gatan K2 summit detector mounted on a Titan Krios set to 300 kV set to counting mode for the data collection (Suloway et al., 2005). The dose rate was ∼4.78 e-/pix/s with frame exposure of 250 ms, with a total exposure time and dose of 13.4 s and 60 e^−^/Å^2^, respectively. MotionCor2 was used for frame alignment and CTF models were determined using GCTF (Zheng et al., 2017). DogPicker was used to pick 666,007 particles, which were then extracted and subsequently 2D-classified in cryoSPARC (Punjani et al., 2017; Voss et al., 2009). Selected 2D classes amounting to 277,453 particles were then fed into the 3D heterogeneous refinement algorithm with 3 classes requested. Of these, only one 3D class (155,394 particles, GSFSC 6.21 Å) amounted to a trimer with Fabs bound and it was advanced to homogenous refinement using C3 symmetry, which resulted in a GSFSC resolution of 4.01 Å. Subsequent non-uniform C3 symmetry refinement resulted in a final GSFSC resolution of 3.87 Å.

#### Model Building and Refinement

The BG505 SOSIP.664 trimer structure from PDB 5ACO was docked into the cryoEM map using UCSF Chimera (Pettersen et al., 2004). Subsequent iterations of manual and Rosetta fragment library based centroid rebuilding and refinement were then performed (Simoncini et al., 2012).The resulting model was then refined using all atom refinement under constraints of the density map. Glycans were manually built in Coot (Emsley et al., 2010).

#### Isolation of monoclonal antibodies

Single cell sorting was performed as previously described (Mason et al., 2016). In brief, rhesus PBMCs were stained with an antibody cocktail of CD3 (clone SP34-2, BD Biosciences), CD4 (clone OKT4, BioLegend), CD8 (clone RPA-T8, BD Biosciences), CD14 (M5E2, BD Biosciences), CD20 (clone 2H7, Biolegend), IgM (clone MHM-88, Biolegend), IgG (clone G18-145, BD Biosciences) and BG505 SOSIP-AviB conjugated with streptavidin-Alexa Fluor 647 (AF647) and streptavidin-Alexa Fluor 488 (AF488) respectively. Cells were stained at room temperature for 20 mins in the dark. Antigen-specific memory B cells (CD3^−^CD4^−^CD8^−^CD14^−^CD20^+^IgM^−^IgG^+^BG505 SOSIP.664 dual positive) were single-cell sorted into 384-well culture plates according to the published protocol (Huang et al., 2013). After 2 weeks of culture, supernatants were harvested and screened for neutralization activity. Ig genes from neutralization positive wells were amplified by RT-PCR, nested PCR, then cloned into expression vectors and expressed in 293F cells.

### DATA AND SOFTWARE AVAILABILITY

3D EM reconstructions have been deposited in the Electron Microscopy Databank (https://www.emdataresource.org/) under the accession numbers listed in the Key Resources Table. The EMD accession numbers of bnAb Fabs used for comparison to polyclonal Fab responses were 5624 (PGT121), 6249 (1NC9), 6250 (CH103), 2625 (8ANC195), 5918 (PGT151), and 8125 (VRC34).

## SUPPLEMENTAL INFORMATION

**Figure S1. Related to Figure 1. 3D sorting and 2D classification of infected naïve animal polyclonal responses.** (A) BZ05 (B) BZ11 (C) BZ13 and (D) BZ15 (E) BZ31, and (F) CB00. 3D sorted classes show variable stoichiometry of binding but do not reveal any new epitopes. While we could not generate 3D reconstructions of two of the naïve animals ((D) and (E)) we were still able to assign epitopes based on comparison with 2D class averages of known comparators. For one animal, CB00, (F) there were no Fabs visible bound to the BG505 trimer.

**Figure S2. Related to Figure 2. 3D sorting of low titer animals.** (A) 12-060 (B) 12-065 (C) 0Q7. EM densities are not shown for animals 12M248, 12-153, and 12-149 because they could not be classified beyond what is shown in Figure 2.

**Figure S3. Related to Figure 3. 3D sorting of high titer animals.** (A) 12-143 (B) 12-137 (C) 12-046. EM densities are not shown for animals 12M169, 11M088, and 409 because they could not be classified beyond what is shown in Figure 3 (D) Negative-stain reconstruction of immunization week 28 serum from animal 12-084 incubated with BG505 SOSIP.664, showing FP-2 and C3/V5-3 responses.

**Figure S4. Related to Figure 4. 3D sorting of high titer animal immunization time points.** (A) 12-137 (B) 11M-088 (C) 4O9, and (D) 12-046. The sample preparation for the animal 12-046 complex was significantly different from the rest of this group as described in Methods Details.

**Figure S5. Related to Figure 5. Sequence alignment of the BG505 SOSIP.664 v5.2 trimer with a subset of the viruses from the 12 virus global panel.** (A) BG505, 398F1 and 25710 show identical amino acid sequences comprising the fusion peptide region (B) negative-stain 3D reconstruction of animal 12-046 complexes with virus 25710. Most likely due to the relatively low neutralization titers the occupancy of the fusion peptide Fabs was very low, resulting in a small number of views and thus poor quality 3D reconstruction. Representative 2D class averages are provided below the corresponding densities.

**Figure S6. Related to Figure 6. 3D map of NHP epitopes (transparent) compared to selected bnAb (solid) sites, showing significant overlap with the polyclonal responses detected in rhesus macaques.**

**Figure S7. Related to Figure 7. Detailed analysis of animal 409 antibody responses.** (A) and (B) Map and docked model of P3C23 showing glycans at positions 448 and 355 facilitating the paratope-epitope interaction (C) CDRH3 resolution of the V1 response (D) and (E) Framework region resolution of the V1 Fab HC (F) Interaction of the V1/V3 Fab with the V3 loop and N332 G) Comparison of PGT121 (PDB: 5CEZ) and 409 V1 polyclonal antibody paratope-epitope interactions, showing PGT121 CDRH3 making contact with GDIR while the 409 V1 loop conformation precluding GDIR contacts with the 409 CDRH3 of V1 polyclonals. (H) The CDRH1 of animal 409 makes contact with a single residue, R327, from the GDIR motif.

